# An integrated cell atlas of the human lung in health and disease

**DOI:** 10.1101/2022.03.10.483747

**Authors:** L Sikkema, D Strobl, L Zappia, E Madissoon, NS Markov, L Zaragosi, M Ansari, M Arguel, L Apperloo, C Bécavin, M Berg, E Chichelnitskiy, M Chung, A Collin, ACA Gay, B Hooshiar Kashani, M Jain, T Kapellos, TM Kole, C Mayr, M von Papen, L Peter, C Ramírez-Suástegui, J Schniering, C Taylor, T Walzthoeni, C Xu, LT Bui, C de Donno, L Dony, M Guo, AJ Gutierrez, L Heumos, N Huang, I Ibarra, N Jackson, P Kadur Lakshminarasimha Murthy, M Lotfollahi, T Tabib, C Talavera-Lopez, K Travaglini, A Wilbrey-Clark, KB Worlock, M Yoshida, Lung Biological Network Consortium, T Desai, O Eickelberg, C Falk, N Kaminski, M Krasnow, R Lafyatis, M Nikolíc, J Powell, J Rajagopal, O Rozenblatt-Rosen, MA Seibold, D Sheppard, D Shepherd, SA Teichmann, A Tsankov, J Whitsett, Y Xu, NE Banovich, P Barbry, TE Duong, KB Meyer, JA Kropski, D Pe’er, HB Schiller, PR Tata, JL Schultze, AV Misharin, MC Nawijn, MD Luecken, F Theis

## Abstract

Organ- and body-scale cell atlases have the potential to transform our understanding of human biology. To capture the variability present in the population, these atlases must include diverse demographics such as age and ethnicity from both healthy and diseased individuals. The growth in both size and number of single-cell datasets, combined with recent advances in computational techniques, for the first time makes it possible to generate such comprehensive large-scale atlases through integration of multiple datasets. Here, we present the integrated Human Lung Cell Atlas (HLCA) combining 46 datasets of the human respiratory system into a single atlas spanning over 2.2 million cells from 444 individuals across health and disease. The HLCA contains a consensus re-annotation of published and newly generated datasets, resolving under- or misannotation of 59% of cells in the original datasets. The HLCA enables recovery of rare cell types, provides consensus marker genes for each cell type, and uncovers gene modules associated with demographic covariates and anatomical location within the respiratory system. To facilitate the use of the HLCA as a reference for single-cell lung research and allow rapid analysis of new data, we provide an interactive web portal to project datasets onto the HLCA. Finally, we demonstrate the value of the HLCA reference for interpreting disease-associated changes. Thus, the HLCA outlines a roadmap for the development and use of organ-scale cell atlases within the Human Cell Atlas.

## INTRODUCTION

Single-cell genomics has been enabling scientists to study tissues at unprecedented scale and resolution^1–3^. Rapid technological improvements over the past decade have allowed datasets to grow both in size and number^4^. This has led consortia, such as the Human Cell Atlas, to pursue the generation of large-scale reference atlases of human organs and the entire human body^5, 6^. Such atlases, capturing the diversity of the cellular landscape of human tissues in health, development, normal aging, lifestyles, environmental exposures, and in disease, will advance insights into the molecular underpinnings of natural variation in healthy tissues, as well as mechanisms of disease inception, progression, remission, and treatment response^7^. To capture diversity on the population scale, these atlases must include data from thousands of individuals. The generation of atlases of this scale by a single group is currently not feasible. However, the cumulative datasets generated by the research community at large now contain hundreds of individual tissue donors, sampled at various anatomical locations within an organ. Integration of these disconnected studies into a single harmonized model may offer superior and much-needed^8^ coverage of relevant donor demographic variables, which are expected to impact the molecular phenotypes of the cells in a tissue.

The power of large-scale single-cell cross-dataset analysis has been demonstrated in several recent studies^9–15^. We previously performed the first single-cell meta-analysis of the respiratory system, aggregating expression data of three genes important for SARS-CoV-2 cell entry from 31 datasets, spanning over 200 individuals^11^. This study showed cell-type specific effects of sex, age and smoking status on gene expression, which could not have been recovered in any individual dataset. Especially when large numbers of individuals, rather than cells, are aggregated, it becomes possible to answer population-level questions using single-cell data. However, currently available integrated atlases are limited in the number of genes^11^, human samples^12, 14^, datasets^14^, or cell types^9, 12^, and include only non-harmonized or limited cell type annotations^9–13^ and subject metadata^9–13^ (e.g. age, BMI, smoking status). These limitations constrain the potential of atlases to serve as a “reference”, as they fail to represent and catalog the diversity of cellular phenotypes within the organ and among individuals. Moreover, choice of integration approach can impact the resulting reference^16^, and only a retinal atlas systematically assessed the quality of data integration^10^.

Overcoming batch effects through data integration is an essential step in building a high-quality atlas and remains one of the key challenges in the field. As each dataset represents a unique view of an organ, containing specific information and biases due to its experimental design, it is paramount to separate technical biases from biologically relevant information when integrating datasets. Successful integration of the available datasets into a single tissue atlas will therefore be a critical step to achieve the goals of the Human Cell Atlas^5^.

Here, we present the first integrated single-cell transcriptomic atlas of the human respiratory system, including the upper and lower airways, from published and newly generated datasets (**fig. 1**). The Human Lung Cell Atlas Core Reference (HLCA core) combines data from 14 datasets, 107 individuals, 168 healthy samples, and 584,000 cells to represent the cellular diversity of the human lung. The HLCA core contains detailed, consensus-based hierarchical cell type annotations with associated marker genes, recovers rare cell types from constituent datasets, and associates cell type-specific gene modules with covariates such as age, BMI, and anatomical location. Through transfer learning, the HLCA core can be used to annotate and interpret new datasets from healthy and diseased individuals, enabling discovery of novel cell types and states associated with normal variability and disease. We demonstrate the power of this approach by extending the HLCA to a total of 2.2 million cells and 444 donors, mapping 32 new human lung datasets from healthy and diseased lungs to the HLCA core. The HLCA thus forms the first consensus single-cell transcriptomic reference for healthy lung tissue.

**Figure 1.**
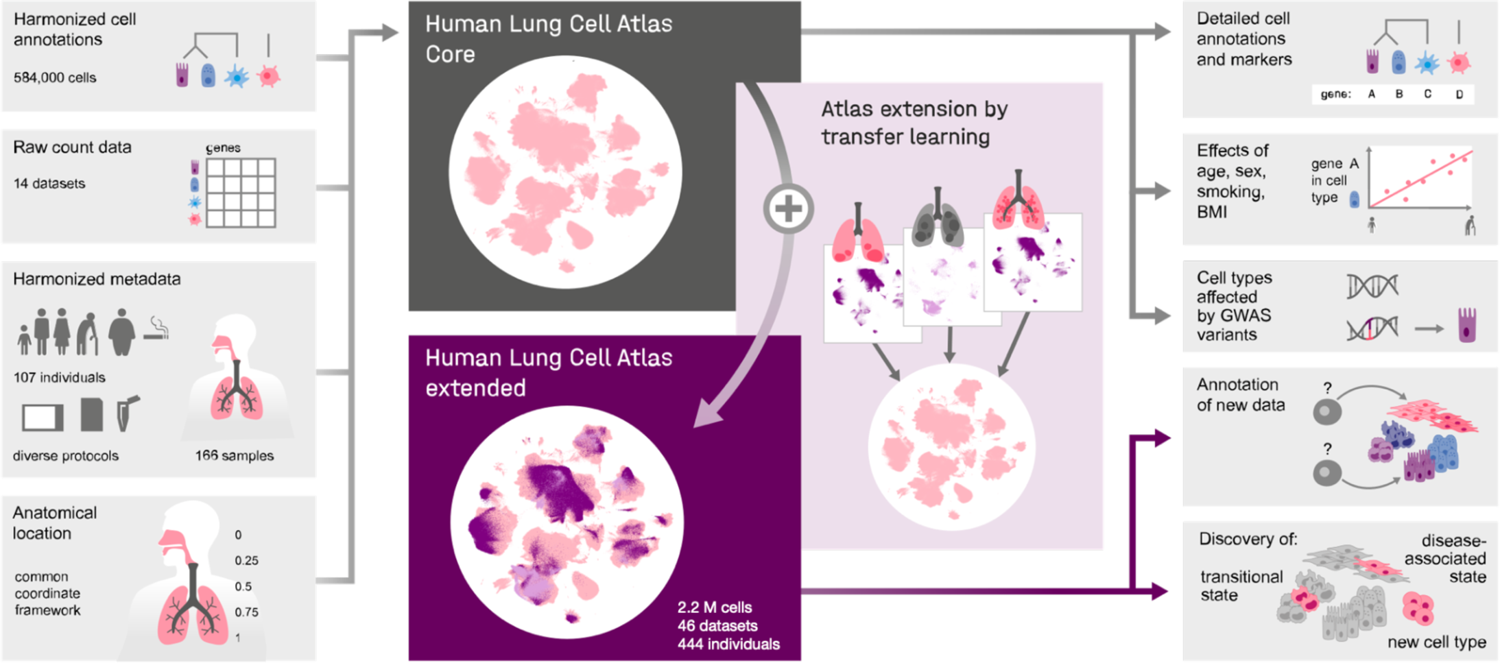
Human Lung Cell Atlas study overview. Harmonized cell annotations, raw count data, harmonized patient and sample metadata, and sample anatomical locations encoded into a common coordinate framework were collected and generated as input for the Human Lung Cell Atlas (HLCA) core (left). After integration of the core datasets, the atlas was extended by mapping 34 additional datasets, including disease samples, to the HLCA core, bringing the total number of cells in the extended HLCA to 2.2 million. The HLCA core provides detailed consensus cell annotations with matched consensus cell type markers (right, top), gene modules associated with technical, demographic, and anatomical covariates in various cellular identities (right, middle), GWAS-based association of lung conditions with cell types (right, middle), a reference projection model to annotate new data (right, middle) and discover new cell types, transitional cell states, and disease-associated cell states (right, bottom).

Together, we provide a roadmap for building and using comprehensive, interpretable, and up-to-date organ- and population-scale cell atlases.

## RESULTS

### Successful data integration into the Human Lung Cell Atlas establishes a core reference

To build a foundation for the HLCA, we collected scRNA-seq data and detailed, harmonized technical, biological and demographic metadata from 14 different datasets (9 published, 5 unpublished)^1, 2, 17–22^. These datasets include samples from 107 individuals, with diversity in age (ranging from age 10 to 76), sex (60% male, 40% female), ethnicity (65% white, 14% black, 2% latino, 2% mixed, 2% Asian, 14% unannotated), BMI (ranging from 20 to 49), and smoking status (52% never, 16% former, 15% active smoker, 17% unannotated) (**fig. 2a**). Cells from the donors were obtained from 166 tissue samples, using a variety of sampling methods, experimental protocols, and sequencing platforms, and coming from different tissue donors (e.g. organ donor, healthy volunteer, **Supplementary Data Table 1, Supplementary Data Table 2**). To obtain a broad reference of the lung and both upper and lower airways, we included samples from nose, lower airways (multiple locations) and lung parenchyma (**Supplementary Data Table 2**). Sample locations were projected onto a 1-dimensional common coordinate framework (CCF) to standardize the anatomical location of origin. The CCF quantifies location along the respiratory tract between the outside world and the alveolar sac in the lung parenchyma by assigning a score between 0 and 1, with 0 representing the most proximal location in the upper airways (inferior turbinate) and 1 representing the most distal location (alveolar sac) (**fig. 2a, Supplementary Data Table 2, Supplementary Data Table 3**).

**Figure 2.**
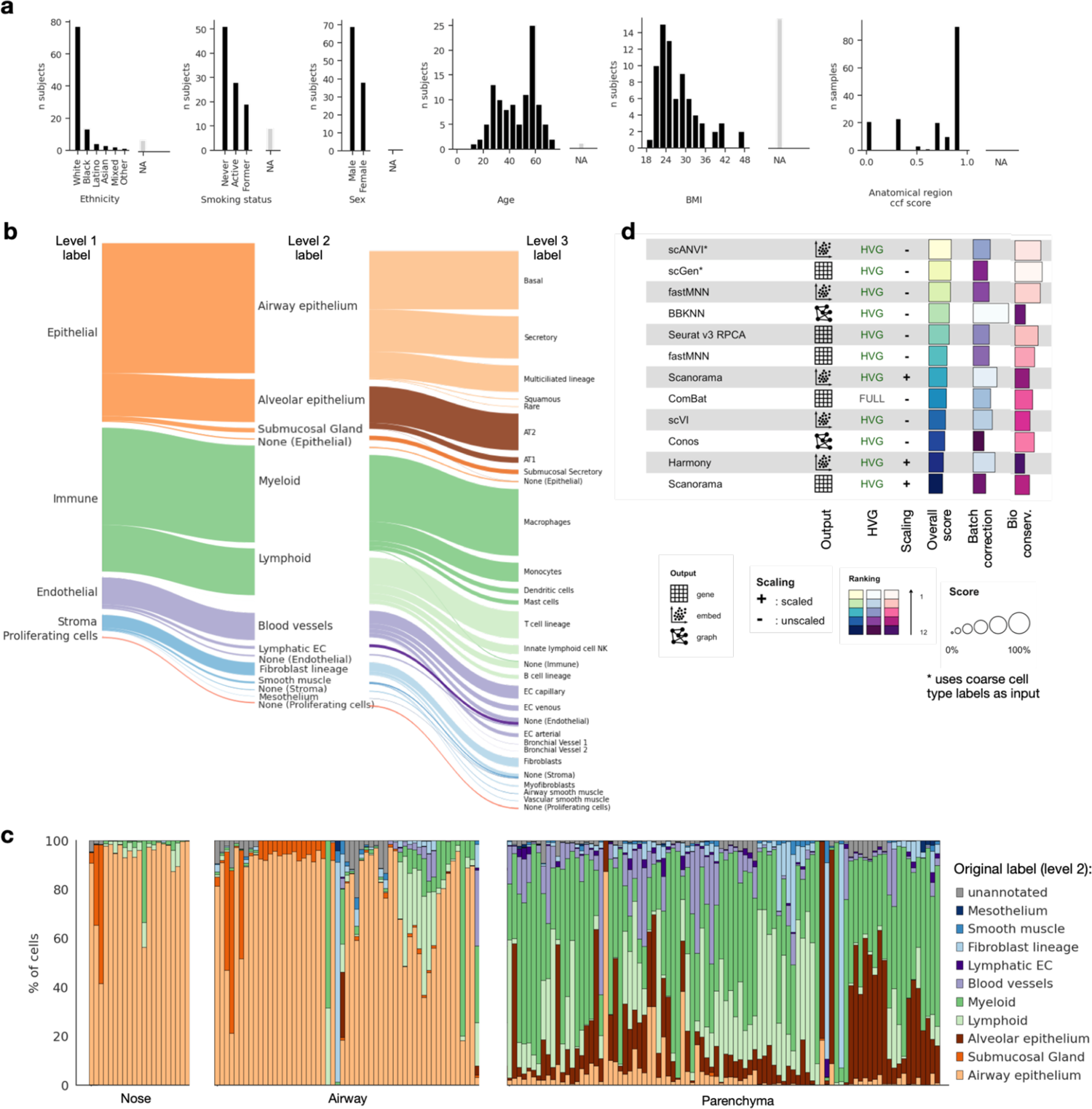
Composition and construction of the HLCA core. **a**, Subject and sample composition in the HLCA core for demographic and anatomical variables, respectively. Subjects/samples without annotation for a specific variable are shown under “NA” (gray bars). Anatomical region common coordinate framework (CCF) score ranges from 0 to 1, with 0 representing the most proximal part of the lung and airways (nose), and 1 the most distal (distal parenchyma). **b**, Overview of the HLCA core cell type composition, based on the 5-level hierarchical reference to which cell type labels of all datasets were mapped. Cell types were labeled up to the finest level of labeling available, and set to “None” if no cell type label was available at the level under consideration. Cell type composition is shown for level 1 (left), 2 (middle) and 3 (right). Level 4 and 5 annotations can be found in **Supplementary Data Table 4**. Each level is broken down into more detailed labels in the following level. Cell labels making up less than 0.02% of all cells are not shown. AT1, 2: alveolar type 1, 2. EC: endothelial cell. NK: natural killer cell. **c**, Cell type composition per sample, based on level 2 labels. Samples are ordered by anatomical region CCF score, and split into nose, airway and parenchyma groups. **d**, Summary of dataset integration benchmarking results. Benchmarking was performed on a subset of 12 datasets from the HLCA. Methods are ordered by overall score. Overall score is a weighted mean of batch-correction and biology-conservation performance. For each method, the results are shown only for the best-performing preprocessing of the data for that method. Preprocessing of data is specified under HVG and scaling. HVG: whether or not genes were subsetted to the 2000 most highly variable genes (“HVG”) or to 6000 genes (“FULL”) before integration. Scaling: whether or not genes were scaled to mean 0 and standard deviation 1 before integration. Output: format of the method’s integrated output.

Single-cell studies in lung tissue have described several novel lung cell types and states^1, 17, 23^. However, consensus definitions of cell types based on single-cell transcriptomic data, particularly of transitional cell states, are lacking. To enable supervised data integration and downstream integrated analysis, we harmonized cell-type nomenclature between the different datasets using a 5-level hierarchical cell identity reference framework (**Supplementary Data Table 4, fig. 2b**). We collected cell identity labels for every dataset as provided by the data generator, and mapped these to the hierarchical reference framework. At the coarsest level (level 1) cells were labeled as epithelial (48%), immune (38%), endothelial (9%), and stromal (4%). Most datasets did not provide the finest level of annotation for all cells, likely due to limitations in cell numbers, such that overall 94%, 66% and 7% of cells were annotated at level 3, 4, and 5, respectively **(fig. 2b**). As expected, cell type composition varied between samples, and especially between locations along the CCF (**fig. 2c**).

To perform integration and remove dataset-specific batch effects, we evaluated 12 different batch integration methods on 12 datasets (**fig. 2d, Extended Data Fig. 1**), using our previously established benchmarking pipeline^24–33^. This benchmarking pipeline includes 12 metrics that evaluate the quality of an integration, quantifying both the removal of batch effects and the conservation of biological variation after integration. Based on these criteria, scANVI^24^ was the top-performing integration method (**fig. 2d**). We therefore integrated the collected datasets using scANVI, creating the HLCA core, a reference atlas of the healthy human respiratory system containing 584,844 cells from 107 individuals (**fig. 3a**).

**Figure 3.**
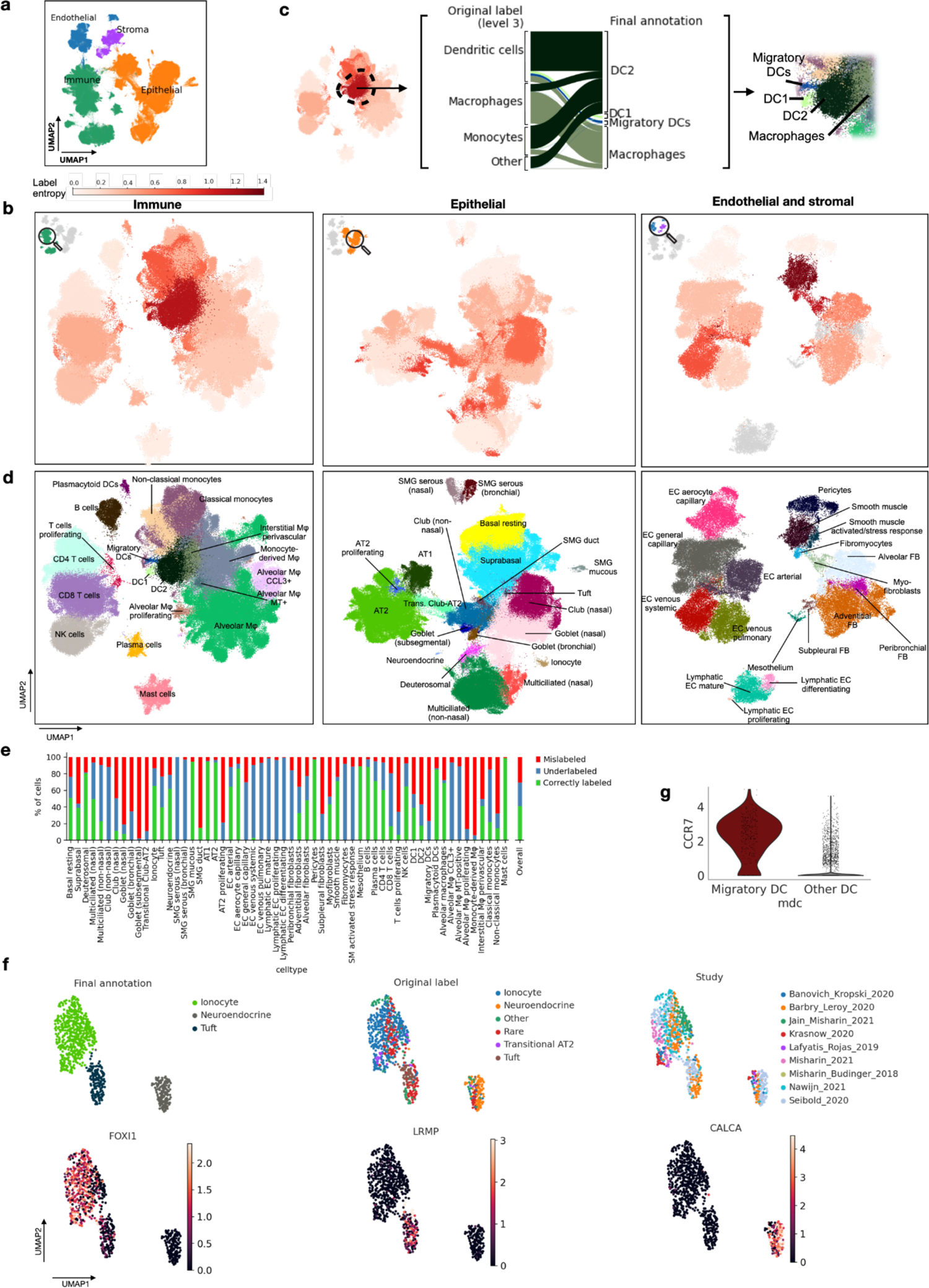
The HLCA core conserves detailed biology and enables consensus-driven annotation. **a**, A Uniform Manifold Approximation and Projection (UMAP) of the integrated HLCA, colored by level-1 annotation. **b**, Cluster label entropy of leiden 3 clusters of the HLCA. HLCA was split into three compartments (immune, epithelial, and endothelial and stromal) for ease of visualization. Cells from every cluster are colored by the shannon-entropy of level 3 labels in the cluster, with higher entropy indicating more contradictory labels in the cluster. Entropy was not calculated for clusters with fewer than 20% of cells annotated at level 3 (clusters colored gray). **c**, Cell type label composition of the immune cluster with the highest cluster entropy (left), with original labels (left), and matching manual re-annotations (right). **d**, UMAPs of the immune, epithelial, and endothelial and stromal compartments of the HLCA core with cell identity annotations from the consensus-based manual re-annotation. **e**, Percentage of cells originally mis-labeled or under-labeled (i.e. only labeled at a coarser level), as compared to final manual re-annotations. Percentages were calculated per manual annotation, as well as across all cells (right bar). **f**, UMAP of HLCA clusters annotated as rare epithelial cell types (i.e. ionocytes, neuroendocrine cells and tuft cells). Final annotations, original labels and study of original are shown (top), as well as expression of ionocyte marker *FOXI1*, tuft cell marker *LRMP*, and neuroendocrine marker *CALCA*. **g**, Log-normalized expression of migratory DC marker *CCR7* in cells identified as migratory DCs versus other DCs. DC: dendritic cell. AT1, 2: alveolar type 1, 2 cells. EC: endothelial cells. FB: fibroblasts. Mφ: macrophages. MT: metallothionein. NK cells: natural killer cells. SM: smooth muscle. SMG: submucosal gland.

### The HLCA includes consensus-based detailed cell identity annotations

A large-scale integrated atlas provides the unique opportunity to systematically investigate the extent of consensus in cell type labels that are found across subjects. To identify areas of consensus and disagreement, we iteratively clustered the HLCA core and investigated subject diversity and cell-type label agreement in these clusters using entropy scores (see **Methods**). Most clusters contained cells from many subjects (median 47 subjects per cluster, range 2-102), as illustrated by high subject entropy (>0.43, **fig. 3b, Extended Data Fig. 2**) for 79 out of 93 final clusters. Low subject entropy clusters (n=14) were largely immune cell clusters (n=13, of which 7 macrophage clusters, 4 T cell clusters and 2 mast cell clusters), representing subject- or group-specific phenotypes. High cell type label entropy can identify both annotation disagreements between studies, and clusters of doublets. Indeed, six small clusters with high label entropy even at the coarsest level of annotation highlighted doublet populations (identified via marker gene expression), not labeled as such in the original datasets. These clusters were removed from the HLCA core, bringing the total number of clusters to 93. Of these 93, 61 clusters showed low label entropy (<0.56), suggesting overall agreement of cell-type labels assigned in the original datasets (**fig. 3a,b**). The remaining 32 clusters exhibited high label entropy, highlighting cellular phenotypes that were differently labeled across datasets (**fig. 3a,b**). For example, one cluster of myeloid cells (cluster 1.2.1) contained cells originally labeled as monocytes, macrophages, and dendritic cells at annotation level 3 (**fig. 3c**). Sub-clustering and marker gene expression suggested that this cluster included a homogeneous subcluster of type 2 dendritic cells (DC2s), and the cells in this subcluster were partly mis-labeled as monocytes and macrophages in the foundational datasets (**fig. 3c, Extended Data Fig. 3**). Similarly, we find several such cases among stromal and epithelial cell populations, as well as in continuous cell phenotypes (e.g., goblet and club cells), for which individual studies set different boundaries between cell type labels. The presence of these high label entropy populations, identifying mislabeled cell types, indicates the need for consensus re-annotation of the integrated atlas. As a first step to achieve such a consensus on the diversity of cell types present in the HLCA core, we performed a full re-annotation on the basis of iterative graph-based clustering of the integrated data. Clusters were annotated taking into account the original cell type labels from each study, and a consensus of opinions from 6 lung experts who evaluated the cluster marker genes, to generate tentative consensus cell identity annotations (**fig. 3d**). While our consensus cell type annotations partly correspond to original labels (**fig. 3e**, e.g. AT1, AT2, mast cells), there were also substantial re-annotations (**fig. 3e**, e.g. submucosal gland (SMG) duct cells, migratory dendritic cells (DCs)) or refinements of the original labels (**fig 3e**, e.g. endothelial cell subtypes). In total, we found that 41% of cells were originally given precise and accurate cell type labels that conform to current consensus knowledge, 28% of cells were correctly labeled but at lower granularity (“under-labeled”), and 31% of cells were originally mis-labeled (**fig. 3e**). Mis-labeled cells often differed from the final annotation at the finest level, but matched at a lower level of granularity (e.g. adventitial instead of alveolar fibroblast, **Extended Data Fig. 4**). Out of 58 final cell types, 15 included mostly mis-labeled cells. A potential reason for this mis-labeling is that marker genes used for the original labeling were based on individual publications, and might not generalize well. Therefore, we established a ‘universal’ set of marker genes that generalizes across lung studies, by calculating cell type marker genes over our 14 datasets for every annotation (**Extended Data Fig. 5, Supplementary Data Table 5**). The fully re-annotated HLCA core thus combines data from a diverse set of studies to provide a carefully curated reference for cell type annotations and marker genes in healthy lung tissue.

**Figure 4.**
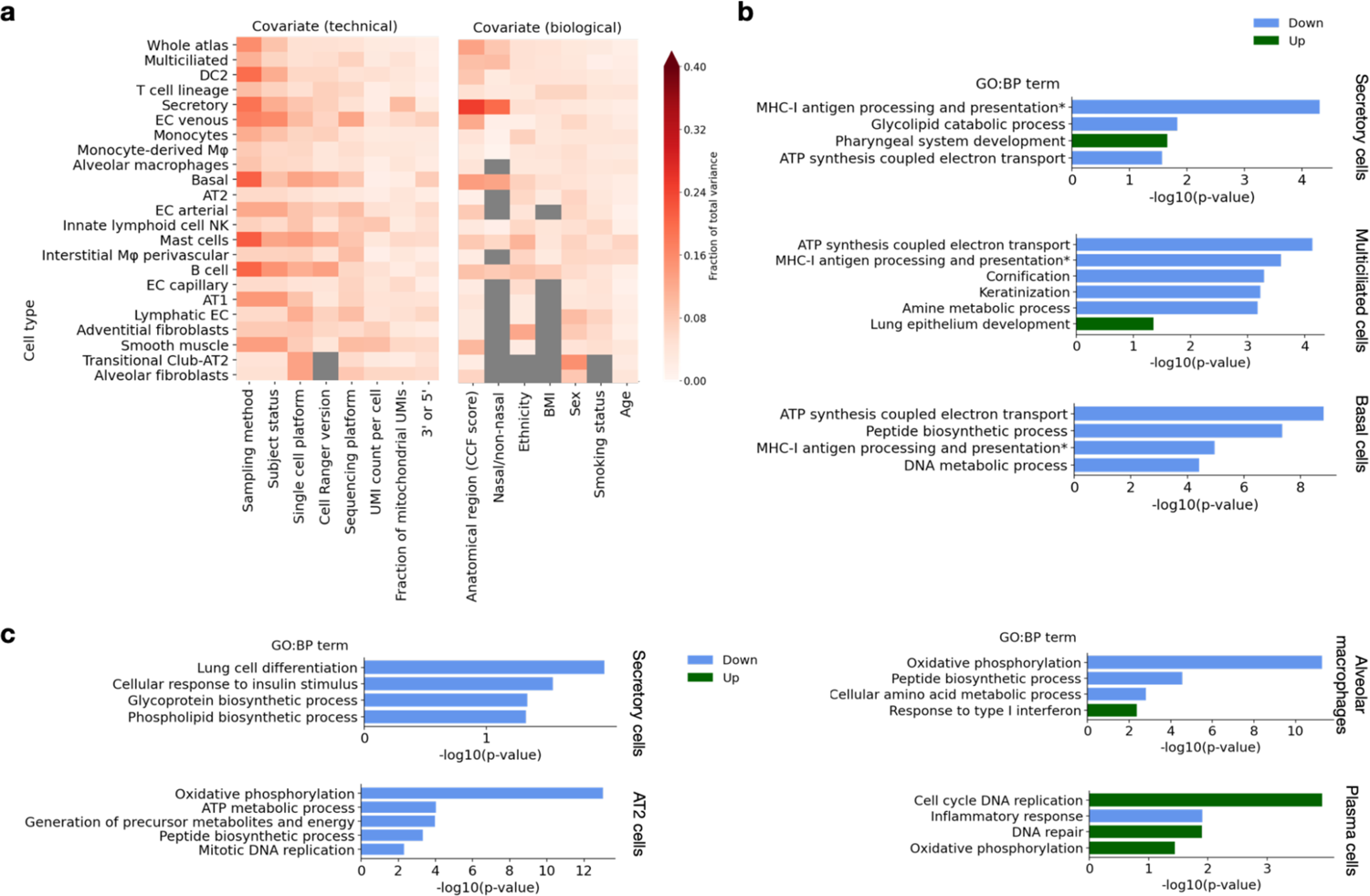
Demographic and technical variables driving inter-individual variation. **a**, Fraction of total inter-sample variance in the HLCA core integrated embedding that can be explained by specific covariates. Covariates are split into technical covariates (left), and biological covariates (right). Cell types are ordered by the number of samples in which they were detected, and only cell types present in at least 40 samples are shown. Fraction of total variance explained is calculated per cell type. Sampling method represents the way a sample was obtained, e.g. surgical resection or nasal brush. Subject status: state of the subject at the moment of sample collection, e.g. organ donor, diseased alive or healthy alive. UMI count per cell: mean of log10(total UMIs detected) among cells of a cell type in a sample. Fraction of mitochondrial UMIs: mean of the fraction of mitochondrial RNA counts across cells from a cell type in a sample. Heatmap is masked gray where fewer than 40 samples were annotated for a specific covariate, or where only one value was observed for all samples for a specific cell type. AT1, 2: Alveolar type 1, 2 cells. EC: endothelial cells. NK: natural killer cells. **b**, Selection of gene sets that are significantly associated with anatomical location, in different epithelial cell types. All gene sets are gene ontology biological process terms. Sets up-regulated towards distal lungs are shown in green, sets down-regulated are shown in blue. *full name: Antigen processing and presentation of peptide antigen via MHC class I. **c**, Selection of gene sets significantly up-(green) or down-regulated (blue) with increasing BMI, in four different cell types. P-values are FDR-corrected using the Benjamini-Hochberg procedure.

**Figure 5.**
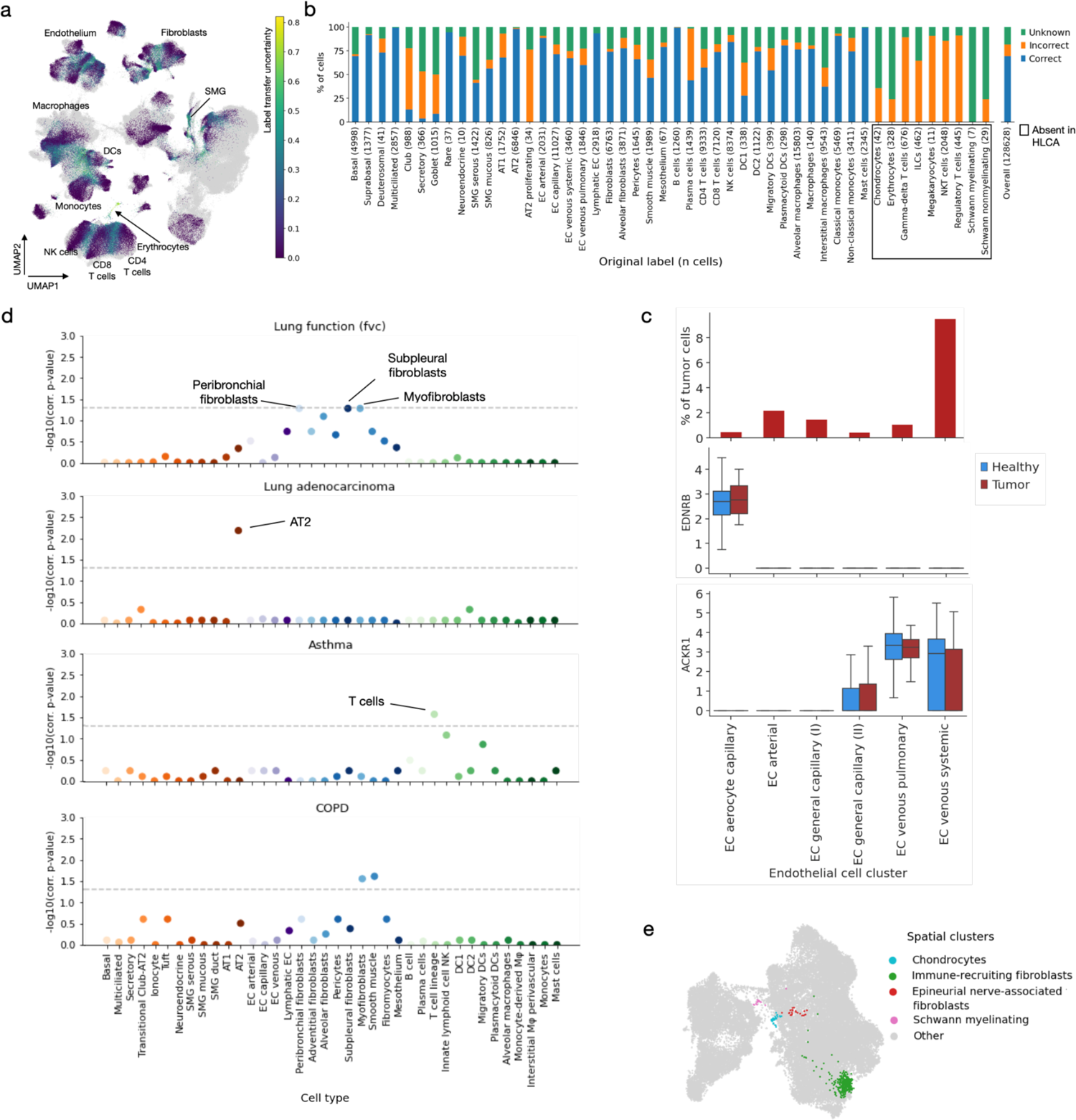
The HLCA core as a reference for label transfer and data contextualization. **a**, UMAP of the jointly embedded HLCA core and the projected healthy lung dataset. UMAP is colored by label transfer uncertainty of cells from the new dataset, with cells from the HLCA colored in gray. HLCA cell types surrounding regions of high uncertainty are labeled. **b**, Percentage of cells from newly mapped healthy lung dataset that are either annotated correctly or incorrectly by label transfer annotation, or annotated as unknown, subdivided by original cell type label. The number of cells in the mapped dataset labeled with each label are shown between brackets after cell type names. Cell type labels not present in the HLCA are boxed. **c**, Top: percentage of cells derived from tumor tissue, per EC cluster from the joint HLCA core and lung cancer data embedding. Only clusters with at least 10 tumor cells are shown. Clusters are named based on the dominant HLCA core cell type annotation in the cluster. Middle: Boxplot of expression of *EDNRB* in EC clusters, subdivided by tissue source. Boxes show median and interquartile range, data points outside more than 1.5 times the interquartile range outside the low and high quartile are considered outliers and not shown. Whiskers extend to the furthest non-outlier point. Bottom: as middle panel, but showing expression of *ACKR1*. **d**, Association of cell types with four different lung phenotypes. Association is based on phenotype-related genomic variants from GWA studies, and cell-type specific differentially expressed genes calculated from the HLCA core. Horizontal dashed lines indicate significance threshold alpha=0.05. p-values are multiple-testing-corrected with the Benjamini-Hochberg procedure. fvc: forced vital capacity. COPD: chronic obstructive pulmonary disease. **e**, UMAP of fibroblast-dominated clusters from the jointly embedded HLCA core and the mapped healthy lung dataset. UMAP is colored by spatial clusters, with cells outside of the indicated clusters colored in gray. AT1, 2: alveolar type 1, 2 cells. DC: dendritic cells. EC: endothelial cells. ILCs: innate lymphoid cells. Mφ: macrophages. NK cells: natural killer cells. NKT cells: natural killer T cells. SMG: submucosal gland.

### The HLCA recovers rare cell types across datasets

Rare cell types such as ionocytes, tuft cells, neuroendocrine cells, or specific immune cell subsets are often hard to identify in individual datasets. To determine whether the higher number of cells in the HLCA core provides better power for identifying rare cell types, we explored the quality of rare cell clustering and their labels. Ionocytes, tuft and neuroendocrine cells make up only 0.08%, 0.01% and 0.02% of the cells in the HLCA core according to the original labels, and were originally identified in only 7, 2 and 4 datasets out of 14, respectively. Despite their low abundance, these cells formed three separate clusters of the HLCA core, with 78%, 76% and 96% of the cells carrying the original label in each cluster (**fig. 3f**). Marker expression shows that the HLCA core correctly excludes cells mis-labeled as rare epithelial cells from these clusters, but also successfully recovers rare cells that were not recognized as such in the original studies (**fig. 3f, Extended Data Fig. 6 a**). Thus, our re-analysis increases the number of datasets in which these rare cells are detected up to three-fold (to 10, 7 and 9 datasets, for ionocytes, tuft and neuroendocrine cells). The correct clustering of rare cell types has been shown to depend on the conservation of subtle biological variation when integrating studies^16^. Importantly, other integration methods tested during our benchmarking, such as Harmony^32^ and Seurat’s RPCA^28^, indeed failed to separate rare cells into distinct clusters, showing the importance of integration method selection when building an atlas (**Extended Data Fig. 6 b**).

**Figure 6.**
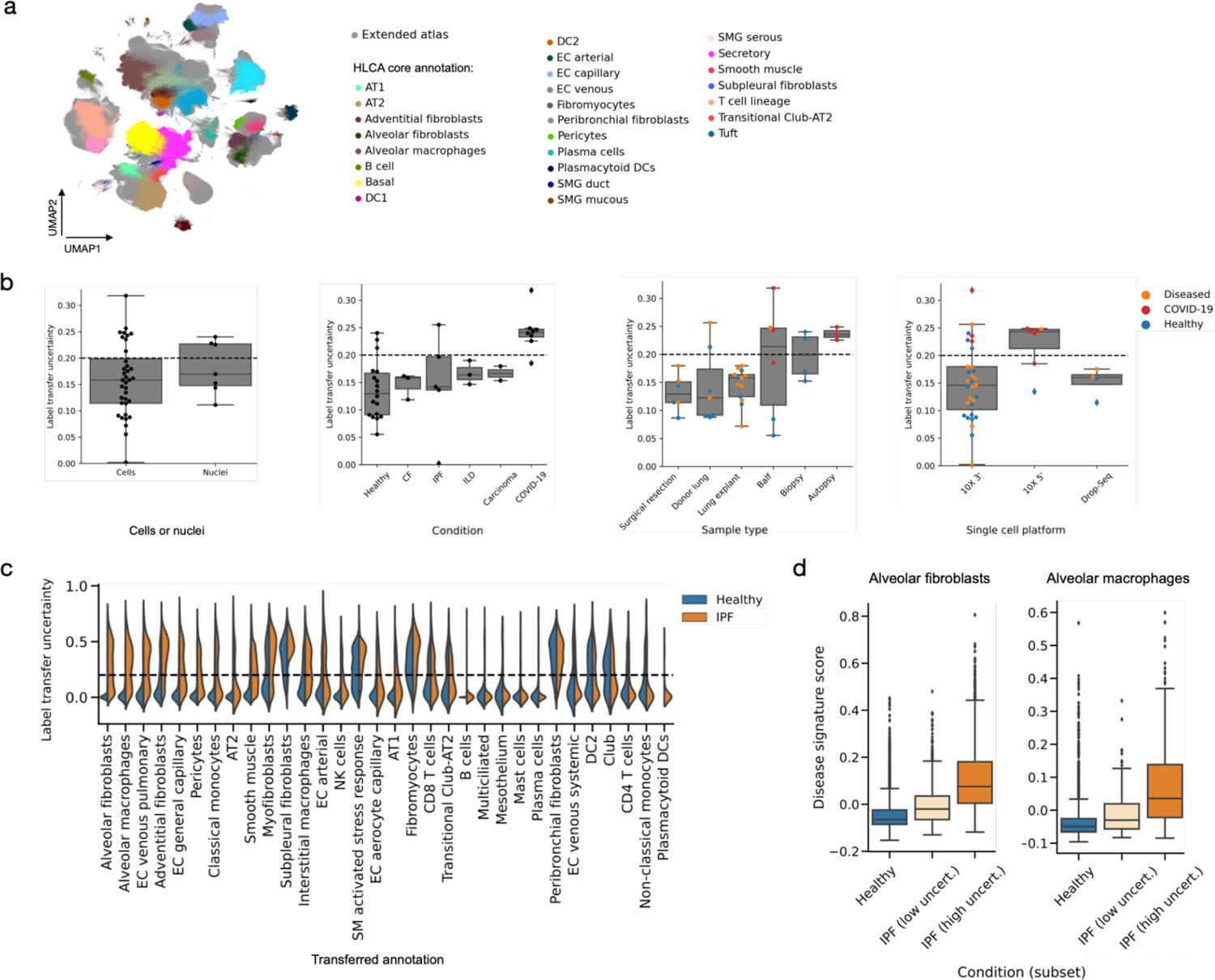
The extended HLCA highlights disease-associated cell states. **a**, UMAP of the HLCA, including both the HLCA core, and the 34 datasets from the atlas extension. Cells from the HLCA core are colored by coarse annotation, whereas cells from the atlas extension (mapped to the HLCA core) are colored gray. **b**, Uncertainty of label transfer from the HLCA core to newly mapped datasets, categorized by several experimental features. Mean uncertainties per dataset are shown. Datasets with samples from multiple categories (e.g. including healthy and diseased samples) were split according to category. Categories with fewer than 2 instances are not shown. CF: cystic fibrosis. IPF: idiopathic pulmonary fibrosis. ILD: interstitial lung disease. **c**, Label transfer uncertainty distribution of cells from a mapped IPF dataset, subdivided by transferred label. Uncertainty is shown for cells from healthy samples (blue) and from IPF samples (orange) from the same dataset. Cell types are sorted by difference in uncertainty between healthy and IPF cells. **d**, Uncertainty-based disease signature scores among alveolar fibroblasts (left) and alveolar macrophages (right), as annotated through label transfer. Cells are split into cells from healthy samples (blue), low-uncertainty cells from IPF samples (light orange), and high-uncertainty cells from IPF samples (bright orange). For both b and d, boxes show median and interquartile range, data points outside more than 1.5 times the interquartile range outside the low and high quartile are considered outliers and not shown. Whiskers extend to the furthest non-outlier point. AT1, 2: alveolar type 1, 2 cells. DC: dendritic cells. DC1, 2: DC type 1, 2. EC: endothelial cells. SMG: submucosal gland.

Due to the increased power in the HLCA core, we were also able to detect rare cell identities that were not labeled in any of the constituent datasets. Through iterative clustering we identified two small clusters (312 cells, 0.05% of all cells) in the immune cell population that contained cells from 12 datasets and thus represent cellular heterogeneity common to multiple studies. Cells from these clusters were a subset of DCs and had distinct and high expression of *CCR7*, a marker for migratory DCs (**fig. 3g, Extended Data Fig. 6 c**)^34, 35^. The HLCA core thus enables improved detection and identification of rare cell types.

### The HLCA reveals cell-type specific effects of technical and demographic variables

Demographic and other biological covariates are known to affect cellular transcriptional phenotypes^11, 22^. Better insight into the impact of these covariates (e.g. sex, BMI, smoking) on cell-type gene expression could shed light on the contribution of the covariates to progression from healthy to diseased states. In addition to these biological covariates, technical covariates such as sample type, or number of transcripts detected per cell can affect transcriptomic variation^16^. The diversity in demographics (e.g. smoking status, age, ethnicity and BMI) and experimental protocols represented in the HLCA core, combined with its single-cell resolution, makes it possible to explore these effects simultaneously and at the level of cell-type specific transcriptomes. To determine the contribution of each covariate to cell-type specific gene expression variation, we calculated the fraction of inter-sample variance that could be explained by the covariate for every cell type (**fig. 4a**, right). To focus on robust correlations detected across studies, we excluded all cell types detected in fewer than 40 samples from this analysis (**Methods, Extended Data Fig. 7a-c**). For many cell types, including epithelial, immune, endothelial and stromal cells, anatomical location is the biological variable explaining most of the variance (**fig. 4a**). In transitional club-AT2 cells interindividual variance is more strongly associated with sex than in any other cell type. Similarly, BMI is most associated with transcriptomic variation in B and T cells, whereas ethnicity is most associated with variation in adventitial fibroblasts. Smoking status is associated with most transcriptomic variation in lymphatic endothelial cells and innate lymphoid/NK cells. Furthermore, for several cell types (e.g. mast cells, basal cells, B cells, and DC2s) the tissue sampling method (brush, biopsy, surgical resection, etc.) explains most variance of all technical as well as biological covariates recorded. Different sampling methods are associated with specific anatomical locations as well as with specific sample processing workflows, and may sample different states of cell activation or differentiation, all of which will contribute to the high degree of variation observed between samples. Nevertheless, despite being sampled with diverse methods, other cell types (e.g. T cells, innate lymphoid/NK cells) are largely unaffected by sampling method (**fig. 4a, Extended Data Fig. 7d**). Therefore, depending on the cell type of interest, sampling methods might need to be harmonized in experimental design. Thus, through this analysis, the HLCA core can help guide important choices in study design.

To better understand how biological variables affect cellular phenotypes, we modeled their cell-type specific effect at the gene level using generalized linear mixed models^36, 37^. We found 936 significant associations of genes with demographic covariates in 29 cell types, and 25,468 significant associations with sample anatomical location (**Supplementary Data Table 6**). Leveraging the diversity in anatomical locations of HLCA core samples, which we encoded into a continuous CCF score (**Supplementary Data Table 3**), we found cell type-specific programs that change with proximal (low CCF score) to distal (high CCF score) location along the respiratory tract. While previous findings highlighted gene expression differences between epithelial cells from the lower airways and the nose^18^, we find that several of these genes also exhibit gradual changes in expression with more distal locations in the tracheobronchial tree. For example, *LMO3*, previously found to be consistently higher expressed in epithelial cells from the nose compared to the lower airways, also displays a gradient along the proximo-distal axis within the tracheobronchial tree. Gene set enrichment analysis shows that gene expression programs that were down-regulated towards distal locations in the tracheobronchial tree in secretory, multiciliated and basal cells were associated with metabolic activity (e.g. cellular respiration, peptide biosynthesis, glycolipid catabolism, **fig. 4b**). Moreover, antigen presentation by MHC-I programs are down-regulated in basal, secretory and multiciliated cells, and keratinization and cornification programs are down-regulated in multiciliated cells towards distal locations along the tracheobronchial tree (**fig. 4b, Supplementary Data Table 7**). In contrast, secretory and multiciliated cells show increased expression of developmental pathway genes along this axis (**fig. 4b, Supplementary Data Table 7**), while basal cells decrease in number (**Extended Data Fig. 8 a**) in accordance with previous findings^38^. These data show presence of gradients in both cell type abundance and transcriptional phenotypes of individual cell types along the tracheobronchial tree, indicating functional adaptations of the cells of the airway wall to their location in the lung.

Similarly, several cell types display transcriptomic changes in subjects with increasing BMI. AT2 cells, secretory cells, plasma cells and alveolar macrophages show the highest numbers of gene sets associated with BMI (**Supplementary Data Table 7**). Of these, AT2 cells, secretory cells and alveolar macrophages exhibit down-regulation of a range of biological processes, including cellular respiration, differentiation, proliferation, and synthesis of peptides and other molecules (**fig. 4c, Extended Data Fig. 8**). In secretory cells, a down-regulation of the insulin-response pathway is also associated with higher BMI, in line with the insulin resistance observed in subjects with obesity^39, 40^ (**fig. 4c**). In alveolar macrophages, inflammatory responses involving JAK/STAT signaling are associated with higher BMI (**fig. 4c, Supplementary Data Table 7**). This signaling pathway has been widely described as a mediator of obesity-induced chronic systemic inflammation^40^, which we can now link to cell type-specific effects in the lung. In contrast to the up-regulation of immune response pathways in alveolar macrophages, in plasma cells high BMI is associated with down-regulation of gene sets associated with immune response, and up-regulation of gene sets associated with cellular respiration, cell cycle and DNA repair (**fig. 4c**). This is consistent with obesity being a known risk factor for multiple myeloma, a plasma cell malignancy^41^. This analysis constitutes a first systematic assessment of the effects of BMI on the lung at single-cell resolution. Thus, the HLCA enables a detailed understanding of the effects of anatomical and demographic covariates on the cellular landscape of the lung, and their relation to disease.

### The HLCA transfers detailed cell-type annotations to new data and identifies unknown cell types

The HLCA core contains an unprecedented diversity of subjects, sampling protocols, and cell identities, and has the potential to serve as a transcriptomic reference for future lung research. Mapping new datasets to this reference can substantially speed up data analysis by generating an informative representation of the data and transferring cell identity annotations. To determine how well annotations can be transferred from the HLCA core to new datasets, we mapped a recently released dataset to the HLCA core using scArches, our recently published transfer learning tool^42^ (**Extended Data Fig. 9, a-c**). This dataset includes several novel cellular identities that were detected by jointly analyzing single-cell, single-nucleus, and spatial data from 10X Visium^43^ to the HLCA core. We transferred cell-type annotations from the HLCA core to the new dataset on the basis of transcriptomic similarity in the batch-corrected joint embedding (**Methods, Extended Data Fig. 9, a-c**). Importantly, the uncertainty of each transferred label (indicating how confidently the label is assigned) is quantified, and cells with high uncertainty scores are labeled as “unknown”. Most cells in the new dataset were annotated with high confidence, indicating a high similarity with the cell types already present in the HLCA core (**fig. 5a, Extended Data Fig. 9d**). High levels of uncertainty were observed specifically in regions representing continuous transitions from one cell type to another, and among cellular identities not currently present in the HLCA core. For example, cells in the continuum between monocytes and dendritic cells displayed high label transfer uncertainty (**fig. 5a**). Several cell identities that were not present in the HLCA core were correctly labeled as “unknown”, even when only few cells of that identity were present in the projected dataset (erythrocytes (n=328), chondrocytes (n=42), Schwann myelinating (n=7), Schwann non-myelinating (n=29); **fig. 5b, Extended Data Fig. 9 e**). Incorrect annotations (i.e. transferred annotations with high certainty, but not matching the original label) highlighted two different cases: cell types that are part of a continuum, with cutoffs between cell types chosen differently in the reference than in the projected data (e.g. secretory, club and goblet cells); and cell types lacking from the HLCA core that have high transcriptional similarity to other cell types that are present in the HLCA, which was observed for several finely annotated immune cell identities. For example, gamma-delta T cells, innate lymphoid cells (ILCs), megakaryocytes, NKT cells, and regulatory T cells were not annotated in the HLCA core, as these cell types could not be distinguished with confidence in the integrated object, and were often lacking in the constituent datasets. Indeed, cell types from the T cell/ILC/NK lineages are known to be particularly difficult to annotate in detail using transcriptomic data only^14^. Therefore, cells annotated with these labels in the projected dataset were largely incorrectly annotated as CD4+ T cells, CD8+ T cells, and NK cells through label transfer (**fig. 5b, Extended Data Fig. 9 e**). Overall, the transferred labels were correct in the majority of cases, with 69% of the cells correctly labeled, 12% of labels incorrect, and 18% set to “unknown”, highlighting the presence of cell types in the projected dataset absent from the HLCA core (**fig. 5b**). Label transfer from the HLCA core thus performs competitively with manual cell type labeling as performed in the original studies (**fig. 3e, 5b**), is highly time-efficient, and provides a higher level of annotation detail. Taken together, the HLCA core can be used as a reference for label transfer to achieve highly detailed annotation of new datasets, and allows identification of new and unknown cell types in those datasets based on label transfer uncertainty. This greatly speeds up the analysis of new lung datasets.

### The HLCA provides crucial context for understanding disease

Single-cell studies of disease rely on adequate, matching control samples to allow correct identification of disease-specific changes. Even when such matching controls are available, it can be challenging to distinguish where disease effects differ from natural variation using a single dataset. The HLCA core can provide this context needed for the study of disease. To demonstrate the ability of the HLCA core to contextualize disease data and facilitate their interpretation, we mapped scRNA-seq data from lung cancer samples^44^ to the HLCA core with scArches (**Extended Data Fig. 10 a-c**). Using HLCA core-based label transfer, we correctly identified cell states missing from the HLCA core as “unknown” (cancer cells and erythroblasts). The remaining cells were annotated correctly in 77%, incorrectly in 1%, and as unknown in 22% of cases (**Extended Data Fig. 10 d-g**). A result of the original study was the separation of endothelial cells into “tumor-associated” and “normal”^44^. Clustering the projected dataset with our reference showed that cells expressing the suggested tumor-associated marker ACKR1 were also abundant in healthy tissue from the HLCA core, specifically in venous ECs (**fig. 5c, Extended Data Fig. 11 a, b**). Indeed, “tumor marker” expression in tumor cells is not higher than that of healthy cells in these clusters (**fig. 5c**). This suggests that ACKR1 is a general marker of venous ECs rather than a tumor-specific EC marker, (**fig. 5c, Extended Data Fig. 11 c**). Similarly, the reported ‘normal EC’ marker EDNRB characterizes aerocyte capillary ECs, both in tumor and in healthy tissue (**fig. 5c, Extended Data Fig. 11 d**). As EC numbers in the original study were low, correctly identifying and distinguishing these cell types without a larger healthy reference is challenging. Thus, without the HLCA core context, limitations in sampling and experimental design can be misinterpreted as meaningful differences between healthy and disease tissue. By mapping disease data to the HLCA core, we can ensure a representative comparison with healthy cells. This projection provides context that is crucial to correctly interpret disease-related phenotypes, especially when healthy controls are limited.

In addition, the HLCA can provide context to the results of large-scale genetic studies of disease. Genome-wide association studies (GWAS) link disease with specific genomic variants that may confer an increased risk of disease. Previous studies have linked such variants to cell-type specific mechanistic hypotheses, which are often lacking in the initial association study^45, 46^. As the HLCA core has a strongly increased resolution in cell type annotations compared to existing atlases^1, 2, 18^, it can serve as a resource for linking disease variants to specific lung cell types at a much higher granularity. To demonstrate the value of the HLCA core in contextualizing genetic data, we mapped association results from four GWA studies of lung function or disease^47–50^ to the HLCA core cell types^51^ (**fig. 5d, Extended Data Fig. 13; Methods**). We show that genomic variants associated with lung function (forced vital capacity) are associated with peribronchial fibroblasts (p=0.052), subpleural fibroblasts (p=0.052), and myofibroblasts (p=0.052), suggesting these fibroblast subtypes play a causative role in inherited differences in lung function (**fig. 5d**). Interestingly, epithelial and immune cell types show no association with lung function. We further find a significant association of lung T cells with asthma-associated single nucleotide polymorphisms (SNPs) (p=0.02). Lung adenocarcinoma-associated variants show a significant association with AT2 cells (p=0.007), and myofibroblasts and smooth muscle cells are significantly associated with COPD GWAS SNPs (p=0.03, p=0.02 resp.) (**fig. 5d**). Thus, by linking genetic predispositions to lung cell types, the HLCA core serves as a valuable resource to improve our understanding of lung function and disease.

### Projecting new data onto the HLCA refines HLCA annotations

As knowledge of cell types in the lung expands, and the size of newly generated datasets increases, annotations in the HLCA core will need to be further refined. The HLCA and its annotations can be updated by learning from new data projected onto the reference. We simulated such an HLCA update using the previously projected healthy lung dataset, specifically focussing on the novel cell identities that were distinguished based on their tissue location in matched spatial transcriptomic data (“spatially annotated cell types”). These cell identities (i.e. immune-recruiting fibroblasts, perichondrial fibroblasts, non-myelinating and myelinating Schwann cells, epineurial and endoneurial nerve-associated fibroblasts, and chondrocytes) were predominantly from the mesenchymal lineage, and were present at very low frequencies (median: 0.004% of all cells, **Extended Data Fig. 12 a**). In the updated atlas, that combines the HLCA core and the projected data, 3 out of 7 spatially annotated cell types (perichondrial fibroblasts, non-myelinating Schwann cells, and endoneurial nerve-associated fibroblasts, 16, 29 and 35 cells, respectively) could not be recovered in distinct clusters, possibly due to low cell numbers in the projected dataset and clustering parameterization (**Methods**). In contrast, all spatially annotated mesenchymal cell types with more than 40 cells (immune-recruiting fibroblasts and chondrocytes), and two rare cell types (myelinating Schwann cells and epineurial nerve-associated fibroblasts), were recovered in distinct clusters (“spatially annotated clusters”) (**fig. 5e, Extended Data Fig. 12 a, b**). Thus, the combined embedding of the HLCA core and the projected dataset retains subtle biological variation present in the newly projected data, which can be used to update the atlas. Moreover, three out of four spatially annotated clusters also contained cells from the HLCA core, thereby enabling a refinement of existing HLCA core annotations using the spatial context from the projected dataset. For example, HLCA core annotations could be extended for a small subset of fibroblasts to immune-recruiting fibroblasts, which we confirmed using marker gene expression (**Extended Data Fig. 12 c**). The fourth spatial cluster (chondrocytes) exclusively contained cells from the projected data, suggesting this cell type was absent in the HLCA but could nonetheless be detected as a separate cluster in the combined embedding. Thus, the HLCA core and its annotations can be refined by mapping new datasets to the atlas and incorporating annotations from these new datasets into the reference.

### The HLCA allows large-scale mapping of new data and identification of new cell types and states in healthy and diseased lungs

To extend the atlas and include samples from lung disease, we mapped 1,647,652 cells from 338 healthy and diseased individuals from 34 datasets (6 unpublished^43^, 28 published^3, 15, 17, 21, 44, 52–66)^ to the HLCA core using scArches^42^, bringing the HLCA to a total of 2.2 million cells from 444 individuals (**fig. 6a**). To test the HLCA as a reference for lung tissue in both health and disease across single-cell technologies, we included healthy samples and samples from numerous lung diseases containing single-cell as well as single-nucleus RNA-seq data, and generated by technologies such as Drop-Seq and SeqWell (**Supplementary Data Table 1**). Label transfer from the HLCA core to the newly mapped datasets enabled detailed cell type annotation across datasets even for rare cells. In comparison, rare cell type identification with conventional annotation approaches^67^ is challenging, especially in smaller datasets, and is linked to several instances of mislabelling (**fig. 3e, Extended Data Fig. 4**). For example, 1,996 migratory DCs could be identified across 25 datasets via label transfer (which constitute 0.1% of all cells in these datasets), whereas this cell type was originally labeled in only 2 of the datasets of the 7 for which cell type labels were available (**Extended Data Fig. 14**).

As label transfer uncertainty highlights cell identities not captured in the HLCA, we investigated whether the uncertainty metric can also be used to identify disease-associated states of healthy cell types captured in the HLCA core. High label transfer uncertainty can indicate (1) insufficient batch removal when mapping new datasets, (2) outlier samples in the mapped data, and (3) cell identities not captured in the HLCA core. Based on correspondence between transferred annotations and original labels of 7 datasets (**Methods**), we found that a mean label transfer uncertainty above 0.2 at the dataset level suggests that residual batch effects (biological or technical) are still present. Out of 34 new datasets, 24 had a mean uncertainty score below this threshold, indicating that mapping of new data to the HLCA core is of high quality in the majority of cases. These 24 datasets include disease samples, as well as single-nucleus and Drop-Seq single-cell data (**fig. 6b**), demonstrating the potential of the HLCA core as a universal reference.

Datasets with a mean uncertainty above 0.2 were often from COVID-19 studies and associated with specific technical covariates (bronchoalveolar lavage fluid (BALF), autopsy, and 10X 5’-sequencing) (**fig. 6b**). As control samples from BALF and 10X 5’ datasets showed uncertainty levels below 0.2, high label transfer uncertainty for the remaining datasets was likely due to disease-related cell identities in the data (not present in the healthy reference), rather than residual technical batch effects. Indeed, high uncertainty values in these datasets were driven by macrophages and monocytes (**Extended Data Fig. 15**), which are activated and recruited in SARS-CoV-2 pneumonia^65, 66^ Pulmonary diseases are characterized by the emergence of unique disease-associated transcriptional phenotypes^2, 17, 19, 21, 68^ We observed higher levels of label transfer uncertainty in datasets from diseased lungs (**fig 6b**, condition), possibly flagging cell types changed in response to disease. For example, labels of various fibroblast and macrophage cell types known to be affected by idiopathic pulmonary fibrosis (IPF) are transferred with higher uncertainty in IPF samples than in samples from healthy controls from the same dataset^62^ (**fig. 6c, Extended Data Fig. 16 a, b**). Label uncertainty thus detects cell identities that are specifically altered in the context of disease, an essential step in the analysis of diseased tissue data.

In addition to detecting transferred annotations altered in disease, we investigated whether uncertainty scores, which are quantified per cell, also directly highlight the cells from these annotations that are most affected by the disease condition, thereby immediately focusing on the relevant populations. To identify these cellular subsets, and learn matching disease-specific gene signatures, we compared gene expression between high-uncertainty and low-uncertainty cells in samples from patients with IPF (**Supplementary Data Table 8, Supplementary Data Table 9**). For both alveolar fibroblasts and macrophages, the genes more highly expressed in high-uncertainty cells are indeed lowly expressed in healthy samples (**fig. 6d**), indicating that uncertainty scores can be used to discover new disease-associated cell states and extract disease-specific gene expression programs. Genes down-regulated in high-uncertainty IPF macrophages compared to their low-uncertainty counterparts are associated with homeostatic functions of tissue-resident alveolar macrophages and lipid metabolism (*PPARG, FABP4,* and others)^19, 21, 57^, while up-regulated genes are associated with extracellular matrix remodeling and scar formation (*SPP1, PLA2G7, CCL2;* **Supplementary Data Table 8**)^19, 21, 57^. A more detailed analysis demonstrated that these differentially expressed genes identify distinct subsets of homeostatic tissue-resident and profibrotic monocyte-derived macrophages (**Extended Data Fig. 16 c**). Thus, the HLCA core can be used to annotate new data, identify previously unreported populations, and detect disease-affected cell states and corresponding gene signatures. This vastly speeds up analysis and interpretation of new data, immediately prioritizing the most relevant populations.

## DISCUSSION

We built the Human Lung Cell Atlas, an integrated reference atlas of the human respiratory system that contains over 2.2 million cells from 46 datasets, covering all major lung single-cell RNA-seq studies published to date. The core of this atlas consists of a fully integrated healthy reference of 14 datasets with a consensus reannotation of, and robust marker genes for 58 cell identities, representing a first data-derived consensus annotation of the cellular landscape of the human lung. We leveraged the unprecedented complexity of the HLCA in the number of cells and samples to optimize recovery of rare cell types which could not be annotated in individual datasets, and to identify cell-type specific gene modules associated with relevant covariates such as lung anatomical location, and age, sex, BMI and smoking status of the tissue donor. To provide the HLCA reference as a tool for the community, we built an scArches reference model of the HLCA that enables efficient projection of new data onto the atlas. By projecting data onto the HLCA, we showed that the HLCA enables a fast, detailed annotation of new datasets, including detection of rare cell identities. When applied to disease data, HLCA reference mapping facilitates direct identification of unique, disease-associated cell states and corresponding gene signatures. Taken together, the HLCA is a universal reference for single-cell lung research that promises to accelerate future studies into pulmonary health and disease.

The constituent datasets of the HLCA widely vary in their experimental design (sampling method, subject selection, single cell platform etc.) and therefore exhibit dataset-specific batch effects. Reduction of such batch effects, while retaining biological information, is crucial for integrated analysis of the data. We showed that the high quality of the HLCA hinged on the choice of integration method, with methods like Seurat’s RPCA^28^ and Harmony^32^ being unable to correctly group rare cell identities into separate clusters. Importantly, integration of datasets inherently entails a trade-off between removal of batch-effect and conservation of biological variation^16^. While conserving biological variation in the HLCA enabled capturing subtle biological heterogeneity, such as rare cell types or altered cell states, it also resulted in some clusters being dominated by cells from single individuals. These clusters may represent residual batch effects or rare biological variation. As we showed that substantial transcriptomic variance is associated with demographic covariates, which vary between individuals, residual differences observed between individuals may be related to biological variation. Most subject- or group-specific clusters are observed in immune cells, which have been reported to show strong subject-specific transcriptome variation^69^. Notwithstanding, we observe gene expression programs that are consistently associated with covariates across the integrated HLCA, indicating that biological variation is well retained.

The ultimate goal of a human lung cell atlas is to provide a comprehensive overview of all cells in the lung, and their variation from individual to individual. The HLCA contains healthy lung samples from 107 individuals in its core, and from 444 individuals in the extended atlas. These sample numbers make it possible to systematically assess the contribution of demographic covariates to inter-individual variation at single-cell resolution for the first time. While the HLCA is the most diverse lung study to date, it is still a reflection of the limited diversity in single-cell studies of the respiratory system. Thus, the HLCA core consists of data from 65% white individuals, highlighting the urgent need for diversification of the population sampled in lung studies. Similarly, the current HLCA core does not include BALF, autopsy or sputum samples. Expanding the atlas with these sample types will further diversify captured cell identities, and could improve the quality of the HLCA core as a reference for new datasets acquired by these sampling methods.

Future extensions of the HLCA could further increase the power to detect the effects of demographic variables, such as age, BMI, and smoking status. Indeed, in the current HLCA the effect of smoking on cell types was relatively modest, also compared to other demographic covariates. As metadata on pack years and smoking frequency was not available for most subjects, a coarser encoding of smoking status (current, ever, never) was necessary, increasing the within-group variance and thus decreasing statistical power. A larger number of subjects in the HLCA would counteract this limitation and could provide the power needed to detect more nuanced effects. Similarly, in the current HLCA core certain subsets of T cells (regulatory T cells, gamma-delta T cells) could not be identified as separate clusters. Mapping of additional, well-annotated immune cell data could provide the power to confidently detect these cell identities. Indeed, we showed that mapping datasets to the HLCA enabled refinement of HLCA annotations.

The HLCA core is publicly available as a data portal and a model repository to explore, download, and use as a reference for new datasets. As the atlassing community has multiple outlets for newly generated data, we made the atlas available in Sfaira^70^, Zenodo^71^, Azimuth^14^, CellTypist^72^, and FASTGenomics^73^. Mapping new datasets to the HLCA core using scArches can be done via interactive portals, such as FASTGenomics and Azimuth, as well as locally using our Zenodo model^74^, which lends itself to integration into bioinformatics pipelines. As more lung datasets become available and the respiratory community progresses in charting the cellular landscape of the lung, for example via novel sampling techniques or multi-modal analysis such as combined spatial and single-cell analysis, knowledge of the lung and its cell types will also expand. This underlines the importance of viewing the HLCA as a ‘live’ resource that requires continuous updates. Indeed, recent studies^1, 17, 22, 43^ have proposed more, novel cell types that have not yet been included in the hierarchical, harmonized HLCA annotations as these do not yet have support from multiple studies or consensus within the field. Using recent advances in transfer learning, we can robustly and iteratively extend the current HLCA. We illustrated, by mapping a newly annotated dataset to the HLCA core, that incorporating novel data in the atlas enables the refinement of HLCA annotations. To consolidate this incorporation of new knowledge, the HLCA core will be continuously updated by adding new datasets as these become available. New builds of the HLCA will be established by fully re-integrating the original and new datasets into an updated HLCA core. In addition, versions of every build will include fast, transfer learning-based mappings of individual, newly published datasets to the latest available HLCA core build. With the integration of new datasets and the open access of all HLCA infrastructure, the HLCA can serve as a community- and data-driven platform for open discussion on lung cell heterogeneity, thus providing the basis for reaching consensus on updated lung cell type annotations. Through this iterative process, the HLCA will function as a central reference for lung data, reflecting the latest knowledge of the lung and its cell identities in health and disease.

In the future, multimodal data of the human respiratory system - including epigenomic, spatial, and imaging data - will become increasingly available. While first ideas for integrating such data have been proposed^75^, a systematic study of multimodal atlas building is not yet feasible due to limited data. Similarly, integrating single-cell atlases with large population cohort studies assaying human transcriptomic variation on the bulk level would greatly extend the represented diversity that is lacking in available single-cell studies. Integration across bulk and single-cell data would however require methodological advances in bulk deconvolution and imputation. Overall, we expect future versions of the HLCA to also contain a multimodal and a spatial “track”, which will enable deeper insights into cellular phenotypes within the organ.

Taken together, the HLCA represents a major milestone for the respiratory community, providing a single-cell resource of unprecedented size and quality. It offers a model framework for building integrated, consensus-based, population-scale atlases for further organs within the Human Cell Atlas. The HLCA is publicly available, and combined with an open-access platform to map new datasets to the atlas, this resource paves the way towards a better and more complete understanding of both health and disease in the human lung.

## METHODS

### Re-processing of published data

The raw sequencing data of 4 previously published studies was re-aligned to GRCh38, ensembl 84 for the HLCA (Krasnow_2020, Barbry_Leroy_2020, Seibold_2020, Meyer_2021). For the Krasnow_2020 and Barbry_Leroy_2020 studies, the CellRanger 3.1.0-based HLCA pipeline was used for re-alignment. Seibold_2020 and Meyer_2021 data were processed as previously described^22, 43^, but with reference genome and genome annotation adapted to the HLCA (GRCh38, ensembl84). All other datasets in the HLCA core were originally already aligned to GRCh38, ensembl84, except data from the Lafyatis_Rojas_2019 study (GRCh38, ensembl93) (**Supplementary Data Table 1**).

### Single-cell sequencing and preprocessing of unpublished data

Unpublished data was generated as follows:

*Barbry_unpubl:* Barbry_unpubl: Nasal and tracheobronchial samples were collected from IPF patients after obtention of their informed consent, following a protocol approved by CHU Nice, registered at clinicaltrials under reference NCT04529993. The libraries were prepared as described in Deprez et al^13^, and yielded an average of 61,000±11,000 cells per sample, with a viability above 95%. The single cell suspension was used to generate single cell libraries following the V3.1 protocol for 3’ chemistry from 10x Genomics (CG000204). Sequencing was performed on a NextSeq 500/550 sequencer (Illumina). Raw sequencing data were processed using the cellranger-6.0.0 pipeline, with reference genome and annotation GRCh38 and ensembl98. For each sample, cells with less than 200 transcripts or more than 40 000 transcripts were filtered out, as well as genes expressed in less than 3 cells. Normalization and log-transformation was done using the standard Scanpy^76^ pipeline. PCA was performed on 1000 HVG to compute 50 PCs and the Louvain algorithm was used for clustering. Those clusters were then annotated by hand for each sample. Raw counts and the thus obtained cell annotations were used as input for the HLCA.

*Schiller_2021:* Tumor-free, uninvolved lung samples (peri-tumor tissues) were obtained during tumor resections at the lung specialist clinic “Asklepios Fachkliniken Munich-Gauting” and accessed through the bioArchive of the Comprehensive Pneumology Center Munich (CPC-M). The study was approved by the local ethics committee of the Ludwig-Maximilians University of Munich, Germany (EK 333-10 and 382-10) and written informed consent was obtained from all patients.

Single-cell suspensions for single-cell RNAseq were generated as previously described^15^. In brief, lung tissue samples were cut into smaller pieces, washed with PBS and enzymatically digested using an enzyme mix composed of dispase, collagenase, elastase and DNAse for 45 min at 37°C while shaking. After inactivating the enzymatic activity with 10% FCS/PBS, dissociated cells were passed through a 70 µm cell strainer, pelleted by centrifugation (300 x g, 5 min), and subjected to red blood cell lysis. After stopping the lysis with 10% FCS/PBS, the cell suspension was passed through a 30 µm strainer and pelleted. Cells were resuspended in 10%FCS/PBS, assessed for viability and counted using a Neubauer hematocytometer. Cell concentration was adjusted to 1,000 cells/µl and around 16,000 cells were loaded on a 10x Genomics Chip G with Chromium Single Cell 3′ v3.1 gel beads and reagents (3′ GEX v3.1, 10x Genomics). Libraries were prepared according to the manufacturer’s protocol (10x Genomics, CG000204_RevD). After quality check, single-cell RNA-seq libraries were pooled and sequenced on a NovaSeq 6000 instrument.

The generation of count matrices was performed by using the Cellranger computational pipeline (v3.1.0, STAR v2.5.3a). The reads were aligned to the hg38 human reference genome (GRCh38, ensembl99). Downstream analysis was performed using the Scanpy^76^ package (v1.8.0). We assessed the quality of our libraries and excluded barcodes with less than 300 genes detected, while retaining those with a number of transcripts between 500 and 30,000. Further, cells with a high proportion (> 15%) of transcript counts derived from mitochondrial-encoded genes were removed. Genes were considered if they were expressed in at least 5 cells. Raw counts of cells that passed filtering were used as input for the HLCA.

*Duong_HuBMAP_unpubl:* All post-mortem human donor lung samples were obtained from the Biorepository for Investigation of Neonatal Diseases of the Lung (BRINDL) supported by the NHLBI LungMAP Human Tissue Core housed at the University of Rochester. Consent, tissue acquisition, and storage protocols can be found on the repository’s website (brindl.urmc.rochester.edu/). Data was collected as part of the Human Biomolecular Atlas Program (HuBMAP). For isolation of single nuclei, 10 cryosections (40 um thickness) from OCT embedded tissue blocks stored at −80℃ were shipped on dry ice and processed according to a published protocol^77^. Single nucleus RNA-sequencing was completed using 10X Chromium Single-Cell 3’ Reagent Kits v3 according to a published protocol^77, 78^. Raw sequencing data was processed using the 10X Cell Ranger v3 pipeline and GRCh38 (hg38) reference genome. For downstream analysis, mitochondrial transcripts and doublets identified by DoubletDetection^79^ v2.4.0 were removed. Samples were then combined and cell barcodes were filtered based on genes detected (>200, <7500) and gene UMI ratio (gene.vs.molecule.cell.filter) using Pagoda2 (<u>github.com/hms-dbmi/pagoda2</u>). Also using Pagoda2 for clustering, counts were normalized to total counts per nucleus. For batch correction, gene expression was scaled to dataset average expression. After variance normalization, all significantly variant genes 4519 were used for PCA. Clustering was done at different k values (50, 100, 200) using the top 50 principal components and the infomap community detection algorithm. Then, principal component and cluster annotations were imported into Seurat^28^ v4.0.0. Differentially expressed genes for all clusters were generated for each k resolution using Seurat FindAllMarkers (only.pos = TRUE, max.cells.per.ident = 1000, logfc.threshold = 0.25, min.pct = 0.25). Clusters were manually annotated based on distinct differentially expressed marker genes. Raw counts and the thus obtained cell annotations were used as input for the HLCA.

*Banovich_Kropski_2020:* Data was a combination of published data^17^ and unpublished data (Vanderbilt IRB nos. 060165 and 171657 and Western IRB no. 20181836). Unpublished data was generated as previously described^17^, using reference genome GRCh38, ensembl84. Both published and unpublished data were combined and processed together using Seurat v4. Cells containing less than 500 identified genes or more than 25% of reads arising from mitochondrial genes were filtered out. Sctransform^80^ with default parameters was performed to normalize and scale the data, PCA was used for dimensionality reduction using the top 3000 most variable genes. Cell type annotation was performed using Seurat^28, 80^ FindTransferAnchors and TransferData functions, using the annotated object from the published data^17^ as reference. Raw counts and the thus obtained cell annotations were used as input for the HLCA.

*Nawijn_2021:* Data was a combination of published^2^ and unpublished data. In both cases, healthy volunteers were recruited for bronchoscopy at the University Medical Center Groningen, after giving informed consent and according to the protocol approved by the Institutional Review Board. Inclusion criteria and tissue processing were performed as previously described^2^. In short, all subjects were 20-65 years old and had a history of smoking <10 pack-years. To exclude respiratory disease, the following criteria were used: absent history of asthma or COPD, no use of asthma or COPD-related medication, a negative provocation test (PC20 methacholine >8 mg/ml), no airflow obstruction (FEV1/forced vital capacity (FVC) ≥ 70%) and absence of lung function impairment (that is FEV1 ≥ 80% predicted). All subjects underwent a bronchoscopy under sedation using a standardized protocol^81^. Nasal brushes were obtained from the lateral inferior turbinate in a subset of the volunteers right before bronchoscopy using Cyto-Pak CytoSoft nasal brush (Medical Packaging Corporation, Camarillo, CA, USA). Six macroscopically adequate endobronchial biopsies were collected for this study, located between the third and sixth generation of the right lower and middle lobe. Bronchial brushes were obtained from a different airway at similar anatomical locations using a CellCellebrity bronchial brush (Boston Scientific, Massachusetts, USA). Extracted biopsies and bronchial and nasal brushes were processed directly, with a maximum of 1 hour delay. Bronchial biopsies were chopped biopsies using a single edge razor blade. A single-cell solution was obtained by tissue digestion using 1mgml−1 collagenase D and 0.1mgml−1 DNase I (Roche) in HBSS (Lonza) at 37 °C for 1h with gentle agitation for both nasal brushes and bronchial biopsies. Single-cell suspensions were filtered forced using 70µm nylon cell strainer (Falcon), followed by centrifugation at 550g, 4 °C for 5min and one wash with a PBS containing 1% BSA (Sigma-Aldrich). The single-cell suspensions used for 10X Genomics scRNA-seq analysis were cleared of red blood cells by using a red blood cell lysis bufer (eBioscience) followed by live cell counting and loading of 10,000 cells per lane. We used 10XGenomics Chromium Single-Cell 3’ Reagent Kits v2 and v3 according to the manufacturers’ instructions. Raw sequencing data were processed using the CellRanger 3.1.0-based HLCA pipeline, with reference genome and annotation GRCh38 and ensembl84. Ambient RNA correction was performed using FastCAR (https://github.com/LungCellAtlas/FastCAR), using an empty library cutoff of 100 UMI and a maximum allowed contamination chance of 0.05, ignoring the mitochondrial RNA. Data were merged and processed using Seurat^28^, filtering to libraries with >500 UMIs and >200 genes and to the libraries containing the lowest 95% of mitochondrial RNA per sample and <25% mitochondrial RNA, normalized using sctransform^80^ while regressing out % mitochondrial RNA. In general, 15 principal components were used for the clustering, at a resolution of 0.5 to facilitate manual annotation of the dataset. Clusters in the final object that were driven by single donors were removed. Raw counts and cell annotations were used as input for the HLCA.

*Jain_Misharin_2021*: Nasal epithelial samples were collected from healthy volunteers who provided informed consent at Northwestern Medicine, Chicago, USA. Protocol was approved by Northwestern University IRB (STU00214826). Briefly, subjects were seated and asked to extend their neck. A nasal curette (Rhino-pro, VWR) was inserted into either nare and gently slid from posterior-to-anterior about one centimeter along the lateral inferior turbinate. Five curettes were obtained per participant. The curette tip was then cut and placed in 2 ml of hypothermosol and stored at 4C until processing. Single cell suspension was generated using cold-active dispase protocol reported by Deprez, Zaragosi and colleagues^18, 82^ with slight modification. Specifically, EDTA was omitted and cells were dispersed by pipetting 20 times every 5 min using a 1ml tip instead of trituration using a 21/23G needle, the final concentration of protease from Bacillus Licheniformis was 10 mg/ml. Total digestion time was 30 min. Following the wash in 4 ml of 0.5% BSA in PBS and centrifugation at 400 rcf for 10 min, cells were resuspended in 0.5% BSA in PBS and counted using Nexcelom K2 Cellometer with AO/PI reagent. This protocol typically yields ∼300-500,000 cells with viability >95%. The resulting single cell suspension was then used to generate single cell libraries following protocol for 5’ V1 (10x Genomics, CG000086 Rev M) or V2 chemistry (10x Genomics, CG000331 Rev A). Excess cells from 2 of the samples were pooled together to generate 1 additional single cell library. After quality check, the libraries were pooled and sequenced on a NovaSeq 6000 instrument. Raw sequencing data were processed using the CellRanger 3.1.0 pipeline, with reference genome and annotation GRCh38 and ensembl84. To assign sample information to cells in the single-cell library prepared from 2 samples, we ran souporcell^83^ v.2.0 for that library and 2 libraries that were prepared from these samples separately. We used common genetic variants prepared by the souporcell authors to separate cells into 2 groups by genotype for each library, and Pearson correlation between the identified genotypes across libraries to establish correspondence between genotype and sample. Cell annotations were assigned to cell clusters based on expert interpretation of marker genes for each cluster. Cell clusters were derived with Seurat^28^ v3.2 workflow: samples were normalized with sctransform^80^, 3000 HVGs selected and integrated, and clusters derived from 30 principal components using Louvain algorithm with default parameters. Clusters with low number of UMIs and high expression of ribosomal or mitochondrial genes were excluded as low-quality. Raw counts and the thus obtained cell annotations were used as input for the HLCA. *Misharin_2020*: Protocol was approved by Northwestern University IRB (STU00212120). Two biopsies that included the main left bronchus and distal parenchyma from the upper lobe were obtained from another donor lung that was not placed for transplant (donor 1). A single biopsy from a distal lung parenchyma (donor 2) was obtained from wedge resection of the donor lung for size reduction during lung transplantation. Lung and airway tissues were infused with a solution of Collagenase D (2 mg/ml) and deoxyribonuclease I (0.1 mg/ml) in RPMI, cut into ∼2-mm pieces, and incubated in 10 ml of digestion buffer with mild agitation for 30 min at 37°C. The resulting single-cell suspension was filtered through a 70-m nylon mesh filter, and digestion was stopped by addition of 10 ml of PBS supplemented with 0.5% BSA and 2 nM EDTA (staining buffer). Cells were pelleted by centrifugation at 300 rcf for 10 min, supernatant was removed, and erythrocytes were lysed using 5 ml of 1× Pharm Lyse solution (BD Pharmingen) for 3 min. The single-cell suspension was resuspended in Fc-Block (Human TruStain FcX, BioLegend) and incubated with CD31 microbeads (Miltenyi Biosciences, 130-091-935), and the positive fraction, containing endothelial cells and macrophages, was collected. The negative fraction was then resuspended in staining buffer, the volume was adjusted so the concentration of cells was always less than 5E6 cells/ml, and the fluorophore-conjugated antibody cocktail was added in 1:1 ratio (EpCAM, Clone 9C4, PE-Cy7, BioLegend, catalog number 324222, RRID:AB_2561506, 1:40; CD206, Clone 19.2, PE, Thermo Fisher Scientific, catalog number 12-2069-42, RRID:AB_10804655, 1:40; CD31, Clone WM59, APC, BioLegend, catalog number 303116, RRID: AB_1877151, 1:40; CD45 Clone HI30, APCCy7, BioLegend, catalog number 304014, RRID: AB_314402, 1:40; HLA-DR, Clone LN3, eFluor450, Thermo Fisher Scientific, catalog number 48-9956-42, RRID:AB_10718248, 1:40). After incubation at 4°C for 30 min, cells were washed with 5 ml of MACS buffer, pelleted by centrifugation, and resuspended in 500 ul of MACS buffer + 2 ul of SYTOX Green viability dye (Thermo Fisher Scientific). Cells were sorted on a FACSAria III SORP instrument using a 100-m nozzle and 20 psi pressure. Macrophages were sorted as live/CD45+HLA-DR+CD206+ cells, epithelial cells were sorted as live/CD45−CD31−EpCAM+, and stromal cells were sorted as live/ CD45−CD31−EpCAM− cells. Cells were sorted into 2% BSA in Dulbecco’s PBS (DPBS, without calcium and magnesium), pelleted by centrifugation at 300 rcf for 5 min at 4°C, and resuspended in 0.1% BSA in DPBS to ∼1000 cells/l concentration. Concentration was confirmed using K2 Cellometer (Nexcelom) with AO/PI reagent, and ∼5000 to 10,000 cells were loaded on a 10x Genomics Chip B with Chromium Single Cell 3′ gel beads and reagents (3′ GEX V3, 10x Genomics). Libraries were prepared according to the manufacturer’s protocol (10x Genomics, CG000183_RevB). After quality check, single-cell RNA-seq libraries were pooled and sequenced on a HiSeq 4000 or NovaSeq 6000 instrument. Raw sequencing data were processed using the CellRanger 3.1.0 pipeline, with reference genome and annotation GRCh38 and ensembl84. Each sample was processed with Seurat^28^ v3.2 workflow to split cells into clusters: data was normalized with sctransform^80^, a number of principal components was selected based on gene weights within the component, and cells were clustered with Louvain algorithm. Clusters with low-quality cells were excluded, while clusters expressing markers of multiple cell types were further sub-clustered by repeating the workflow only on those cells. Cell annotations were assigned to clusters based on marker genes for each cluster. Raw counts and the thus obtained cell annotations were used as input for the HLCA.

### Data and metadata collection

To accommodate data protection legislation, single cell RNA sequencing datasets of healthy lung tissue were shared by dataset generators (**Supplementary Data Table 2**) as raw count matrices, thereby obviating the need to share genetic information. Unpublished raw count matrices and cell annotations were generated as described above. For the Barbry_2020 study, count matrices provided had ambient RNA removed. Count matrices were generated using varying software (**Supplementary Data Table 2**). All count matrices of the HLCA core except one study (see above, “re-processing of published data”) were based on alignment to reference genome GRCh38, annotation ensembl 84. For all datasets from the HLCA core, a pre-formatted sample metadata form was filled out by the dataset providers for every sample, containing metadata such as subject ID of the donor from which the sample came, the donor’s age, the type of sample, the sequencing platform, etc. (**Supplementary Data Table 2**). Cell type annotations from dataset providers were included in all datasets.

### Cell type reference creation and metadata harmonization for the HLCA core

To harmonize cell type labels from different datasets, a common reference was created to which original cell type labels were mapped (**Supplementary Data Table 4**). To accommodate labels at different levels of detail, the cell type reference was made hierarchical, with level 1 containing the coarsest possible labels (immune, epithelial, etc.), and level 5 containing the finest possible labels (e.g. naive CD4 T cells). Levels were created in a data-driven fashion, recursively breaking up coarser level labels into finer ones where finer labels were available. Anatomical location of the sample was encoded into a continuous score, with 0 representing the most proximal samples (nose, inferior turbinate) and 1 representing the most distal possible sample (parenchyma, alveolar sacs) (**Supplementary Data Table 3**). A distinction was made between upper and lower airways. First, an anatomical coordinate score was applied to the upper airways, starting at 0 and increasing linearly (with a value of 0.5) between each of the following anatomical locations: inferior turbinate, nasopharynx, oropharnyx, vesibula, and larynx. The trachea received the next anatomical coordinate score using the same linear increment as in the upper airways (score of 2.5). In the lower airways, the coordinate score within the bronchial tree was based on the generation airway, with trachea being the first generation and the total number of generations assumed to be 23^84^. The alveolar sac was assigned the coordinate score of the 23rd generation airway. The coordinate score of each generation airway was calculated by taking the log-2 value of the generation, and adding that to the score of the trachea. Using this methodology, the alveolus received the anatomical coordinate score of 7.02. To calculate the finat CCF coordinate, the coordinates scores (ranging from 0 to 7.02) were scaled to a value between 0 (inferior turbinate) and 1 (alveolus). Samples were then mapped to this coordinate system using the best approximation of the sampling location for each of the samples of the Core HLCA.

### General data preprocessing for the HLCA core

Patients with lung conditions affecting larger parts of the lung, such as asthma or pulmonary fibrosis, were excluded from the HLCA core, and later added to the extended atlas. For the HLCA core, all matrices were gene-filtered based on cellranger ensembl84 gene type filtering^85^ (resulting in 33694 gene IDs). Cells with fewer than 200 genes detected were removed (removing 2335 cells, and 21 extra erythrocytes with close to 200 genes expressed, these hampered SCRAN normalization, see below), and genes expressed in fewer than 10 cells were removed (removing 5167 out of 33694 genes).

### Total counts normalization with SCRAN

To normalize for differences in total unique molecular identifier (UMI) counts per cell, we performed SCRAN normalization^86^. Since SCRAN assumes that at least half of the genes in the data to normalize are not differentially expressed between subgroups of cells, we performed SCRAN normalization within clusters. To that end, we first performed a total counts normalization, by dividing each count by its cell’s total counts, and multiplying by 10,000. We then performed a log transformation using natural log and pseudocount 1. A principal component analysis was subsequently performed. Using the first 50 principal components, a neighborhood graph was calculated with the number of neighbors set to k=15. Data were subsequently clustered with louvain clustering, at resolution r=0.5. SCRAN normalization was then performed on the raw counts, using the louvain clusters as input clusters, and with the minimum mean (library size-adjusted) average count of genes to be used for normalization set to 0.1. The resulting size factors were used for normalization. For the final HLCA (and not the benchmarking subset), cells with abnormally low size factors (<0.01) or abnormally high total counts after normalization (>10e5) were removed from the data (267 cells in total).

### Preprocessing of data for integration benchmarking

For computational efficiency, benchmarking was performed on a subset of the total atlas, including data from 10 studies split into 13 datasets (Lafyatis_Rojas_2020 was split into 10Xv1 and 10Xv2 data, Seibold_2020 was split into 10Xv2 and 10Xv3 data, and Banovich_Kropski_2020 was split into two based on processing site). The data came from 72 subjects, 124 samples and 372,111 cells. Preprocessing of the benchmarking data included the filtering of cells (minimum number of total UMI counts: 500) and genes (minimum number of cells expressing the gene: 5).

For integration benchmarking, the scIB benchmarking framework was used^16^. All benchmarked methods except scGen (i.e. BBKNN, ComBat, Conos, fastMNN, Harmony, Scanorama, scANVI, scVI and Seurat RPCA) were run at least twice: on the 2000 most highly variable genes, and on the 6000 most highly variable genes. Of those methods, all methods that did not require raw counts as input were run twice on each gene set: once with gene counts scaled to have mean 0 and standard deviation 1, one with unscaled gene counts. scVI and scANVI, which require raw counts as input, were not run on scaled gene counts. scGen was only tested on 2000 unscaled highly variable genes. For highly variable gene selection, first, highly variable genes were calculated per dataset, using cellranger-based highly variable gene selection^87^ (default parameter settings: min_disp=0.5, min_mean=0.0125, max_mean=3, span=0.3, n_bins=20). Then, genes that were highly variable in all datasets were considered overall highly variable, followed by genes highly variable in all datasets but one, in all datasets but two etc. until a predetermined number of genes was selected (2000 or 6000 genes).

For scANVI and scVI, genes were subset to the highly variable gene set and the resulting raw count matrix was used as input. For all other methods, SCRAN-normalized (as described above) data were used. Genes were then subset to the pre-calculated highly variable gene sets. For integration of gene-scaled data, all genes were scaled to have mean 0 and standard deviation 1.

Two integration methods allowed for input of cell type labels to guide the integration: scGen and scANVI. As labels, level 3 harmonized cell type labels were used (**Supplementary Data Table 4**), except for blood vessel endothelial, fibroblast lineage, mesothelial and smooth muscle cells, for which we used level 2 labels. Since scGen does not accept unlabeled cells, cells that did not have annotations available at these levels (i.e. cells annotated as cycling, epithelial, stromal, or lymphoid cells with no further annotations; 4499 cells in total) were left out of the benchmarking data.

### Data integration benchmarking

Integration methods were run using default integration parameter settings as previously described^42^. Maximum memory usage was set to 376Gb, and all methods requiring more memory were excluded from the analysis. The quality of each of the integrations was scored using 12 metrics, with 4 metrics measuring the batch correction quality, and 8 metrics quantifying conservation of biological signal after integration (**Extended Data Fig. 1**, metrics previously described^16^). Overall scores were computed by taking a 40:60 weighted mean of batch effect removal to biological variation conservation (bio-conservation), respectively. Methods were ranked based on overall score (**Extended Data Fig. 1**). The top performing method (scANVI, 2000 hvgs, unscaled) was used for integration of the HLCA core.

### Splitting of studies into datasets

For integration of the data into the HLCA core, we first determined in which cases studies had to be split into separate datasets (which were treated as batches during integration). Reasons for possible splitting were 1) different 10X versions used within a study (e.g. 10Xv2 versus 10Xv3), or 2) processing of samples at different institutes within a study. To determine if these covariates caused batch effects within a study, we performed principal component regression^88^. To that end, we preprocessed single studies separately (total counts normalization to median total counts across cells, and subsequent principal component analysis (PCA) with 50 principal components (PCs). For each study, we then calculated the fraction of the variance in the first 50 PCs that could be explained (“pcexpl”) by the covariate of interest (i.e. 10X version or processing institute):

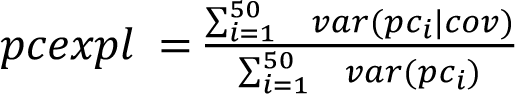

where var(pcI|cov) is the variance in scores for the ith PC across cells that can be explained by the covariate under consideration, based on a linear regression.

Then, 10X version or processing institute assignments were randomly shuffled between samples, and pcexpl was calculated for the randomized covariate. This was repeated over 10 random shufflings, and the mean and standard deviation of the pcexpl was then calculated for the covariate. If the non-randomized pcexpl was more than 1.5 standard deviations above the randomized pcexpl, we considered the covariate a source of batch effect and split the study into separate datasets. Thus both Jain_Misharin_2021 and Lafyatis_Rojas_2019 were split into 10Xv1 and 10Xv2, Seibold_2020 was split into 10Xv2 and 10Xv3, while Banovich_Kropski_2020 was not split based on 10X version nor based on processing location.

### Dataset integration

For integration of the datasets into the HLCA core, coarse cell type labels were used as described for integration benchmarking, except cells with lacking annotations were set to “unlabeled” instead of being removed. scANVI was run on the raw counts of the 2000 most highly variable genes (calculated as described above), using datasets as the batch variable. The following parameter settings were used: number of layers: 2, number of latent dimensions: 30, encode covariates: True, deeply inject covariates: False, use layer norm: both, use batch norm: none, gene likelihood: nb, n epochs unsupervised: 500, n epochs semi-supervised: 200, frequency: 1. For the unsupervised training, the following early-stopping parameters were used: early stopping metric: elbo, save best state metric: elbo, patience: 10, threshold: 0, reduce lr on plateau: True, lr patience: 8, lr_factor: 0.1. For the semi-supervised training, the following early-stopping-parameter settings were used: early stopping metric: accuracy, save best state metric: accuracy, on: full dataset, patience: 10, threshold: 0.001, reduce lr on plateau: True, lr_patience: 8, lr_factor: 0.1. The latent embedding generated by scANVI was used for downstream analysis (clustering and visualization). For gene-level analyses (differential expression, covariate effect modeling) un-corrected counts were used.

### Data embedding and clustering

To cluster the cells in the HLCA core, a nearest neighbor graph was calculated, based on the 30 latent dimensions that were obtained from the scANVI output, with the number of neighbors set to k=30. This choice of k, while improving clustering robustness, could impair the detection of very rare cell types. Coarse leiden clustering was done on the graph with resolution r=0.01. For each of the resulting level 1 clusters, a new neighbor graph was calculated using scANVIs 30 latent dimensions, with the number of neighbors again set to k=30. Based on the new neighbor graph, each cluster was clustered into smaller “level 2” clusters with leiden clustering at resolution r=0.2. The same was done for level 3, 4 and where needed 5, with k set to 15, 10, and 10 respectively, and resolution set to 0.2. Clusters were named based on their parent clusters and sister clusters, e.g. cluster 1.2 is the third biggest subcluster (starting at 0) of cluster 1. For visualization, a two-dimensional Uniform Manifold Approximation and Projection^89^ (UMAP) of the atlas was generated based on the 30-nearest-neighbor graph.

### Calculation of cluster entropy of cell type labels and subjects

To calculate cluster cell type label entropy for a specific level of annotation, Shannon entropy was calculated as 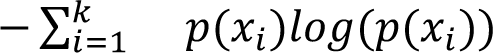, where *x_1_… x_k_* are the set of labels at that annotation level, and *p(x_i_)* is the fraction of cells in the cluster that was labeled as *x_i_*. Cells without a label at the level under consideration were not included in the entropy calculation. If fewer than 20% of cells were labeled at the level, entropy was set to NA. Entropy of subjects per cluster was calculated in the same way. To set a threshold for “high label entropy”, we calculated the label entropy of a hypothetical cluster with 75% of cells given one label, and 25% of cells given another label. Clusters with a label entropy above that level (0.56) were considered high label entropy clusters. To set a threshold for “low subject entropy”, we calculated the label entropy for a hypothetical cluster with 95% of cells from one subject, and the remaining 5% of cells distributed over all other subjects. Clusters with a subject entropy below that level (0.43) were considered clusters with low subject entropy.

### Rare cell type analysis

To determine how well rare cell types (ionocytes, neuroendocrine cells and tuft cells) were clustered together and separate from other cell types after integration, we calculated recall (% of all cells annotated as a specific rare cell type that were grouped into the cluster) and precision (% of cells from the cluster that were annotated as a specific rare cell type) for all level 3 clusters. Nested clustering on Harmony^32, 89^ and Seurat’s RPCA^28^ output was done based on PCA of the corrected gene counts, re-calculating the PCs for every parent cluster when performing clustering into smaller children clusters, and proceeding clustering as described above. The level 3 clusters with the highest sensitivity for each cell type are shown in figures.

### Manual cell type annotation and marker gene selection

Re-annotation of cells in the HLCA core was done by six investigators with expertise in lung biology (EM, MCN, AVM, LEZ, NEB, JAK), based on original annotations and differentially expressed genes of the HLCA core clusters. Annotation was done per cluster, using finer clusters where these represented specific known cell types or states rather than subject-specific variation. Annotations were hierarchical (as the cell type reference), and each annotated cluster was annotated at all levels up to its highest known level.

Marker genes were calculated by performing a t-test-based differential expression analysis on a given cell type (based on SCRAN-normalized, logarithmized counts), compared to the rest of the cells from the same level 1 annotation (i.e. epithelial, endothelial, stromal or immune cells). Genes were then filtered further according to the following criteria: minimum fraction of cells within the cluster in which the gene was detected: 0.25; maximum fraction of cells from outside the cluster in which gene was detected; 0.5, minimum fold change; 1. The top 10 most significant genes that passed filtering were selected as top marker genes. For highly similar cell types, a second differential expression analysis was done, this time comparing the cell type to its closest sister annotations (e.g. submucosal gland (SMG) serous (nasal) cells compared to the remaining SMG cell types). For those cell types, top 10 marker genes consisted of the top 5 genes from the first and second comparison, respectively. Where fewer than 10 genes passed filtering criteria, only those genes were included as markers.

### Variance between individuals explained by technical and biological covariates

To quantify to what extent different technical and biological covariates correlated with inter-individual variation in the atlas, we calculated the “variance explained” by each covariate for each cell type. We first split the data in the HLCA core by cell type annotation, merging substates of a single cell type into one (**Supplementary Data Table 10**). For every cell type, we excluded samples that had fewer than 10 cells of the sample. We then summarized covariate values per sample for every cell type as follows: mean across cells from sample for scANVI latent components (integration results), UMI count per cell, and fraction of mitochondrial UMIs; for all other covariates (i.e. dataset, 3’ or 5’, BMI, cell ranger version (1, 2, 3 or 4), cell viability %, ethnicity, sampling method (e.g. biopsy, brush, donor lung), sequencing platform, sex, single cell chemistry (e.g. 10X 3’ v2), smoking status (active, former or never), subject status (alive disease, alive healthy or organ donor), age, and anatomical region (CCF score)), each sample had only one value, therefore these values were used.

We then performed “PC regression” on every covariate, as described earlier, now using scANVI latent component scores instead of principal component scores for the regression. Samples that did not have a value for a given covariate (e.g. where BMI was not recorded for the subject) were excluded from the regression. Categorical covariates were converted to dummy variables. Celltype - covariate pairs for which only one value was observed for the covariate were excluded from the analysis.

To check to what extent covariates correlated with each other, thereby possibly acting as confounders in the PC regression scores, we determined dependence between all covariate pairs for every cell type. If at least one covariate was continuous, we calculated the fraction of variance in the continuous covariate that could be explained by the other covariate (dummying categorical covariates), and took the square root (equal to Pearson r for two continuous covariates). For two categorical covariates, if both covariates had more than two unique values we calculated normalized mutual information (NMI) between the covariates using scikit-learn^90^, since a linear regression between these two covariates is not possible.

To control for spurious correlations between inter-individual cell type variation and covariates due to low sample numbers, we assessed the relationship between sample number and mean variance explained across all covariates for every cell type. We found that for cell types sampled in fewer than 40 samples, mean variance explained across all covariates showed a high negative correlation with the number of samples (**Extended Data Fig. 7, a**). We reasoned that for these cell types, correlations between interindividual variation and our covariates were inflated due to under-sampling. Moreover, we note that at lower sample numbers, technical and biological covariates often strongly correlate with each other across subjects (**Extended Data Fig. 7, c**). This might lead to the attribution of true biological variation to technical covariates, and vice versa, complicating interpretation of observed inter-individual cell type variation. Consequently, we consider 40 a recommended minimum number of samples to avoid spurious correlations between observed inter-individual variation and tested covariated, and excluded results from cell types with fewer samples.

### Modeling variation between individuals at the gene level and gene set enrichment analysis

To model the effect of demographic and anatomical covariates (sex, age, BMI, ethnicity, smoking status, and anatomical location of sample) on gene expression, we employed a generalized linear mixed model (GLMM). We used sample-level pseudo-bulks (split by cell type), since the covariates modeled also varied at sample- or subject level, and not at cell level. Modeling these covariates at cell level, i.e. treating single cells as independent samples even when coming from the same sample, has been shown to inflate p-values^36, 37^. We encoded smoking status as a continuous covariate, setting never to 0, former to 0.5, and current to 1. Anatomical region was encoded into anatomical region ccf-scores as described earlier. As we noted that changes from nose to the rest of the airways and lungs were often independent from continuous changes observed in the lungs only, we encoded “nasal” as a separate covariate, setting samples from the nose to 1, and all others to 0. BMI and age were re-scaled, such that the 10th percentile of observed values across the atlas was set to 0, and the 90th percentile set to 1 (25 and 64 for age, respectively, and 21.32 and 36,86 for BMI). First, we split the lung cell atlas by cell type annotation, pooling detailed annotations into one subtype (e.g. grouping all lymphatic EC subtypes into one) (**Supplementary Data Table 10**). We then proceeded the same way for every cell type: we filtered out all genes that were expressed in fewer than 50 cells, and all samples that had fewer than 10 cells of the cell type. We furthermore filtered out datasets with fewer than 2 subjects, and refrained from modeling categories in covariates that had fewer than three subjects in their category, for that cell type. To determine if covariance between covariates was low enough for modeling, we calculated the variance inflation factor (VIF) between covariates at subject level. The VIF quantifies multi-collinearity among covariates of an ordinary least squares regression, and a high VIF indicates strong linear dependence between variables. If the VIF was higher than 5 for any covariate for a specific cell type, we concluded covariance was too high and excluded that cell type from the modeling. As many cell types lacked sufficient representation of other ethnicities than ‘’white”, while including ethnicity in the analysis simultaneously reduced the samples that could be included in the analysis to only those with ethnicity annotations, we excluded ethnicity from the modeling.

Gene counts were summed across cells for every sample, within cell type. Sample-wise sums (i.e. pseudo-bulks) were normalized using edgeR’s calcNormFactors function, using default parameter settings. We then used voom^91^, a method designed for bulk RNA-seq that estimates observation-specific gene variances and incorporates these into the modeling. Specifically, we used a voom extension (differential expression testing with linear mixed models or “Dream”) that allows for mixed effect modeling, and modeled gene expression as:

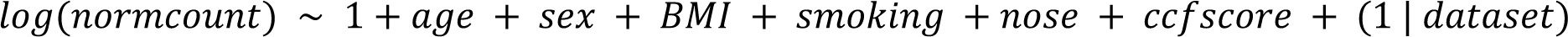

where dataset is treated as a random effect, and all other effects are modeled as fixed effects. Resulting p-values were corrected for multiple testing within every covariate using the Benjamini-Hochberg procedure.

Gene set enrichment analysis was performed in R^92^ using the cameraPR function^93^ in the limma package^94^ with the differential expression test statistic. Gene ontology (GO) biological process terms^95, 96^ were tested separately for each comparison. These sets were obtained from MSigDB (v7.1)^97^ as provided by the Walter and Eliza Hall Institute (https://bioinf.wehi.edu.au/MSigDB/index.html).

### Mapping of GWAS results to the lung cell atlas cell types

To stratify GWAS results from several lung diseases by lung cell type, we applied stratified sc-LDSC, a method that can link GWAS results to cell types based on proximity of disease-associated variants to genes differentially expressed in the cell type^51^. GWAS summary statistics of COPD^49^ (GWAS catalog ID: GCST007692, dbGaP accession number: phs000179.v6.p2, n_cases=35,735, n_controls=222,076), and of lung cancer^48^ (GWAS catalog ID: GCST004748, dbGaP accession number: phs001273.v3.p2, n_cases=29,266, n_controls=56,450) were made available on dbGap upon request. Summary statistics of lung function^50^ (GWAS catalog ID: GCST007429, n=321,047 individuals), of asthma^47^ (GWAS catalog ID: GCST010043, n_cases=88,486 n_controls=447,859), and of depression^98^ (used as negative control, GWAS catalog ID: GCST005902, n_cases=113,769, n_controls=208,811) were publicly available. A differential gene expression test was done for every grouped cell type (**Supplementary Data Table 10**) in the HLCA using a t-test, testing against the rest of the atlas. The top 1000 most significant genes with positive fold-changes were stored as genes characterizing that cell type (“cell type genes”), and used as input for sc-LDSC. Gene coordinates of cell type genes were obtained based on the GRCh37.13 genome annotation. For single nucleotide-polymorphism (SNP) data (names, locations, and linkage-related information), the 1000 genomes European reference (GRCh37) was used, as previously described^51^. Only SNPs from the HapMap3 project were included in the analysis. For identification of SNPs in the vicinity of cell type genes, we used a window size of 100,000 base pairs. Genes from X and Y chromosomes, as well as HLA genes, were excluded because of their unusual genetic architecture and linkage patterns. For ld-score calculation, a 1 centiMorgan window was used. P-values yielded by scLDSC were corrected for multiple testing for every disease tested, using the Benjamini-Hochberg correction procedure. As a negative control, the analysis was performed with a GWAS of depression, and no cell types were found to be significant (**Extended Data Fig. 13**).

### Extension of the HLCA core by mapping of scRNA-seq and snRNA-seq lung data

To map unseen scRNA-seq and snRNA-seq data to the HLCA, we used scArches, our transfer-learning based method that enables mapping of new data to an existing reference atlas^42^. scArches trains an adaptor added to a reference embedding model, thereby enabling it to generate a common embedding of the new data and the reference, allowing re-analysis and *de novo* clustering of the joint data. The data to map was subsetted to the same 2000 highly variable genes (“hvgs”) that were used for HLCA integration and embedding, and hvgs that were absent in the new data were set to 0 counts for all cells. Raw counts were used as input for scArches. Healthy lung data (Meyer_2021) was split into two datasets: 3’ and 5’-based. Lung cancer data (Thienpont_2018) was also split into two datasets: 10Xv1 and 10Xv2.

The model that was learned previously for HLCA integration using scANVI was used as the basis for the scArches mapping. scArches was then run to train adapter weights that allowed for mapping of new data into the existing HLCA embedding, using the following parameters: freeze-dropout: true, surgery_epochs: 500, train base model: false, metrics to monitor: accuracy and elbo, weight-decay: 0, frequency: 1. The following early-stopping criteria were used: early stopping metric: elbo, save best state metric: elbo, on: full dataset, patience: 10, threshold: 0.001, reduce lr on plateau: True, lr patience: 8, lr_factor: 0.1.

### Identification of clusters with spatially annotated cell types

The Meyer_2021 study of healthy lung included cell type annotations based on matched spatial transcriptomic data. Many of these annotations were not present in the HLCA core. To determine if these new “spatial cell types” could still be recovered after mapping to the HLCA core, we looked for clusters specifically grouping these cells. We focused on seven spatial cell types: perichondrial fibroblasts, epineurial nerve-associated fibroblasts, endoneurial nerve-associated fibroblasts, CCL adventitial fibroblasts, chondrocytes, myelinating schwann cells, and non-myelinating schwann cells. As these cell types were often present at very small frequencies, we performed clustering at different resolutions to determine if these cells were clustered separately at any of these resolutions. We clustered at resolutions 0.1, 0.2, 0.5, 1, 2, 3, 5, 10, 15, 20, 25, 30, 50, 80, and 100. Minimum recall (% of cells with the spatial cell type annotation captured in cluster) was set to 20%, and minimum precision (% of cells from Meyer_2021 study in the cluster that had the spatial cell type annotation) was set to 25%. The cluster with the highest recall was selected for every spatial cell type (unless this cluster decreased precision by more than 50% compared to the cluster with the second highest recall). If precision of the next best cluster was doubled compared to the cluster with highest recall, and recall did not decrease with more than 20%, this cluster was selected.

### Cell type label transfer from the HLCA core to new datasets

To perform label transfer from the HLCA core to the mapped datasets from the extended HLCA, we used the scArches k-nearest neighbor based label transfer algorithm^42^. Briefly, a k-nearest neighbor graph was generated from the joint embedding of the HLCA core and the new, mapped dataset, setting the number of neighbors to k=50. Based on the abundance in a cell’s neighborhood of reference cells of different types, the most likely cell type label for that cell was selected, and a matching uncertainty score was calculated. For label transfer to lung cancer and healthy, spatially annotated projected data (**fig. 5b, e**), cells with an uncertainty score above 0.3 were set to “Unknown”. For the extended atlas, we calibrated the uncertainty score cutoff by determining which uncertainty levels indicate possible failure of label transfer. To determine the uncertainty score at which technical variability from residual batch effects impairs correct label transfer, we evaluated how label transfer performed at the level of datasets, as these predominantly differ in experimental design. To determine an uncertainty threshold indicative of possible failure of label transfer, we harmonized original labels for 7 projected datasets^17, 21, 53, 57, 62, 64^ (one unpublished: “Duong_lungMAP_unpubl”) and assessed the correspondence between original labels with the transferred annotations. To assess the optimal uncertainty cutoff for labeling a new cell as “unknown”, we used these results to generate an ROC curve. We chose a cutoff around the elbow point, keeping the false positive rate below 0.5 (uncertainty cutoff 0.2, true positive rate 0.88, false positive rate 0.49) to best distinguish correct from incorrect label transfers (**Extended Data Fig. 17**). False positives are either due to incorrect label transfer, or due to incorrect annotations in the original datasets. Cells with an uncertainty higher than 0.2 were set to “unknown”.

### Disease signature score calculation

To learn disease-specific signatures based on label-transfer uncertainty scores, cells from the mapped data with the same transferred label were split into low-uncertainty cells (<0.2) and high-uncertainty cells (>0.4). We then performed a differential expression analysis on SCRAN-normalized counts using diffxpy^99^ with default parameters, comparing high- and low-uncertainty cells, including only genes that were expressed in at least 20 cells. The 20 most up-regulated genes based on log-fold changes were selected, after filtering out genes with an FDR-corrected p-value above 0.05 and genes with a mean expression below 0.1 in the high-uncertainty group. To calculate the “score” of a cell for the given set of genes, the average expression of the set of genes was calculated, after which the average expression of a reference set of genes was subtracted from the original average, as previously described^100^. The reference set consists of a randomly sampled set of genes for each binned expression value. The resulting score was considered the cell’s “disease signature score”.

### Version information

diffxpy: 0.7.4+18.gb8c6ae0

edgeR: 3.28.1, R 4.1.1 (covariate modeling)

LDSC: 1.0.1

Limma: 3.46.0, R 4.0.3 (GSEA)

Scanpy: 1.7.2

scArches: 0.3.5

scIB: 0.1.1

scikit-learn: 0.24.1

scvi-tools (scANVI): 0.8.1

### Code availability

The HLCA pipeline for processing of sequencing data to count matrices, used for a subset of HLCA datasets (**Methods**): https://github.com/LungCellAtlas/scRNAseq_pipelines. Code used for HLCA project: https://github.com/LungCellAtlas/HLCA_reproducibility Code for users to map new data to the HLCA core (for automated mapping, see HLCA platforms below): https://github.com/LungCellAtlas/mapping_data_to_the_HLCA

### HLCA data portals and mapping to the HLCA

Automated mapping to the HLCA and label transfer can be done with scArches^42^ at FASTGenomics (https://beta.fastgenomics.org/p/hlca)

Automated mapping to the HLCA, and label transfer with Azimuth^14, 42^ (not shown in manuscript) can be done at: azimuth.hubmapconsortium.org.

Label transfer with CellTypist^72^ (not shown in manuscript): https://www.celltypist.org/models

## Data availability

The HLCA (raw and normalized counts, integrated embedding, cell type annotations and clinical and technical metadata) is publicly available and can be downloaded via FASTGenomics: https://beta.fastgenomics.org/p/hlca or via cellxgene https://cellxgene.cziscience.com/collections/6f6d381a-7701-4781-935c-db10d30de293 The HLCA core reference model and embedding for local mapping to the HLCA can moreover be found on Zenodo: https://zenodo.org/record/6337966#.Yid5Vi9Q28U.

The original, published datasets that were included in the HLCA can be accessed under GEO accession numbers GSE135893, GSE143868, GSE128033, GSE121611, GSE134174, GSE150674, GSE151928, GSE136831, GSE128169, GSE171668, GSE132771, GSE126030, GSE161382, GSE155249, GSE135851, GSE145926, EGA study IDs EGAS00001004082, EGAS00001004344, EGAD00001005064, EGAD00001005065, and under urls https://www.synapse.org/#!Synapse:syn21041850,

https://data.humancellatlas.org/explore/projects/c4077b3c-5c98-4d26-a614-246d12c2e5d7, https://www.ncbi.nlm.nih.gov/projects/gap/cgi-bin/study.cgi?study_id=phs001750.v1.p1, https://www.nupulmonary.org/covid-19-ms2/?ds=full&meta=SampleName, https://figshare.com/articles/dataset/Single-cell_RNA-Seq_of_human_primary_lung_and_bronchial_epithelium_cells/11981034/1, https://covid19.lambrechtslab.org/downloads/Allcells.counts.rds, https://s3.amazonaws.com/dp-lab-data-public/lung-development-cancer-progression/PATIENT_LUNG_ADENOCARCINOMA_ANNOTATED.h5, https://github.com/theislab/2020_Mayr, https://static-content.springer.com/esm/art%3A10.1038%2Fs41586-018-0449-8/MediaObjects/41586_2018_449_MOESM4_ESM.zip, http://blueprint.lambrechtslab.org/#/099de49a-cd68-4db1-82c1-cc7acd3c6d14/*/welcome (see also **Supplementary Data Table 1**).

## Author contributions

LS integrated data and analyzed integrated data. AVM, FJT, LS, MCN, MDL wrote manuscript. AVM, FJT, MCN, MDL supervised analysis of integrated data. AAH, BHK, CHM, CJT, CM, JS, LA, LBW, LEZ, MB, MJA, MVB, TMK, YC generated unpublished data. AVM, CF, HBS, JAK, JLS, KBM, MCN, MJ, NEB, PB, PRT, SAT, SL, TED supervised unpublished data generation and analysis. AC, ACAG, CB, CHM, LEZ, MA, MB, NSM, PKLM analyzed unpublished data. DCS performed integration benchmarking. LZ performed GSEA. AVM, EM, JAK, LEZ, MCN, NEB annotated integrated data. AW, CB, EM, KBW, KT, LB, LBW, MG, MY, NJ, PKLM, TT gathered metadata. AG, KT, NH, NJ re-aligned published data. CD, CRS, CTL, DCS, ILI, LD, LH, LS, LZ curated data. CB, LS, NSM, TW set up a shareable CellRanger pipeline. ML provided scArches support. LS, MP set up an automated scArches mapping pipeline. CX set up CellTypist automated annotation. AAH, AC, ACAG, AG, AMT, AVM, AW, BHK, CB, CD, CF, CHM, CJT, CM, CRS, CS, CTL, CX, DCS, DP, DPS, DS, EC, EM, FJT, GP, HBS, ILI, JAK, JEP, JL, JLS, JR, JS, JSH, JW, KBM, KBW, KT, KZ, LA, LB, LBW, LD, LEZ, LH, LP, LS, LZ, MA, MAS, MB, MC, MCN, MDL, MG, MJ, MJA, MK, ML, MN, MP, MVB, MY, MZN, NEB, NH, NJ, NK, NSM, OE, OR, PB, PH, PKLM, PRT, RL, SAT, SL, TED, TJD, TK, TMK, TT, TW, XS, YC, YX reviewed manuscript.

## Group authors

Lung Biological network group authors: M van den Berge, Y Chen, JS Hagood, AA Hassan, P Horvath, J Lundeberg, S Leroy, C Marquette, G Pryhuber, C Samakovlis, X Sun, LB Ware, K Zhang.

## Competing interests

PRT serves as a consultant for Surrozen Inc., Cellarity Inc., and Celldom Inc. and currently acting CEO of Iolux Inc. FJT consults for Immunai Inc., Singularity Bio B.V., CytoReason Ltd, and Omniscope Ltd, and has ownership interest in Dermagnostix GmbH and Cellarity. JAK reports grants/contracts from Boehringer Ingleheim, Bristol-Myers-Squibb, consulting fees from Janssen and Boehringer Ingelheim, study support from Genentech, and is a member of the scientific advisory board of APIE Therapeutics. SAT, in the past three years, has received remuneration for consulting and Scientific Advisory Board Membership from Genentech, Roche, Biogen, GlaxoSmithKline, Foresite Labs and Qiagen. SAT is a co-founder, board member and holds equity in Transition Bio. DS is a founder of Pliant Therapeutics, a member of the Genentech Scientific Advisory Board and has a sponsored research agreement with Abbvie. OE serves in advisory capacity to Pieris Pharmaceuticals, Blade Therapeutics, Delta 4 and YAP Therapeutics. NK served as a consultant to Boehringer Ingelheim, Third Rock, Pliant, Samumed, NuMedii, Theravance, LifeMax, Three Lake Partners, Optikira, Astra Zeneca, RohBar, Veracyte, Augmanity, CSL Behring, Galapagos, and Thyron over the last 3 years, reports Equity in Pliant and Thyron, and a grant from Veracyte, Boehringer Ingelheim, BMS and non-financial support from MiRagen and Astra Zeneca. NK has IP on novel biomarkers and therapeutics in IPF licensed to Biotech. OR is a co-inventor on patent applications filed by the Broad Institute for inventions related to single cell genomics. ORR is an employee of Genentech since October 19, 2020.

## Funding

This work was supported by:

The National Institute of Health R01HL145372 (JAK/NEB). The Fondation pour la Recherche Médicale (DEQ20180339158), the Conseil Départemental des Alpes Maritimes (2016-294DGADSH-CV), the Inserm Cross-cutting Scientific Program HuDeCA 2018, ANR SAHARRA (ANR-19-CE14–0027), ANR-19-P3IA-0002– 3IA, the National Infrastructure France Génomique (ANR-10-INBS-09-03), H2020-SC1-BHC-2018-2020 Discovair (grant agreement 874656) and the Chan Zuckerberg Initiative, LLC Seed Network grant (CZF2019-002438) “Lung Cell Atlas 1.0”. NIH 1U54HL145608-01. Wellcome (WT211276/Z/18/Z and Sanger core grant WT206194); ESPOD fellowship of EMBL-EBI and Sanger Institute. R01 HL153312; U19 AI135964; P01 AG049665. The Netherlands Lung Foundation project nos. 5.1.14.020 and 4.1.18.226. The National Institute of Health R01HL146557 and R01HL153375. The Helmholtz Association’s Initiative and Networking Fund through Helmholtz AI [ZT-I-PF-5-01]. The Bavarian Ministry of Science and the Arts in the framework of the Bavarian Research Association “ForInter” (Interaction of human brain cells). The National Institute of Health R01HL145372 (JAK/NEB) and the Doris Duke Charitable Foundation (JAK). The Joachim Herz Foundation. The Ministry of Economic Affairs and Climate Policy by means of the PPP. 3IA Cote d’auzr PhD program. R01 HL135156, R01 MD010443, R01 HL128439, P01 HL132821, P01 HL107202, R01 HL117004, and DOD Grant W81WH-16-2-0018. HL142568 and HL14507 from the NHLBI. P50 AR060780-06A1. MRC Clinician Scientist Fellowship (MR/W00111X/1). The Jikei University School of Medicine. University College London, Birkbeck MRC Doctoral Training Programme. CZI and 5U01HL148856; CZI; 5U01HL148856; R01 HL153045; R01HL127349, R01HL141852, U01HL145567. 2R01HL068702, 5R01HL14254903, 4UH3CA25513503, R21HL156124, R56HL157632, R21HL161760. NIH U54 AG075931, 5R01 HL146519. Swedish Research Council Vr, Cancerfonden. CZI Deep Visual Proteomics. 1U54HL145608-01, U01HL148861-03.

## SUPPLEMENTARY FIGURES

**Extended Data Figure 1.**
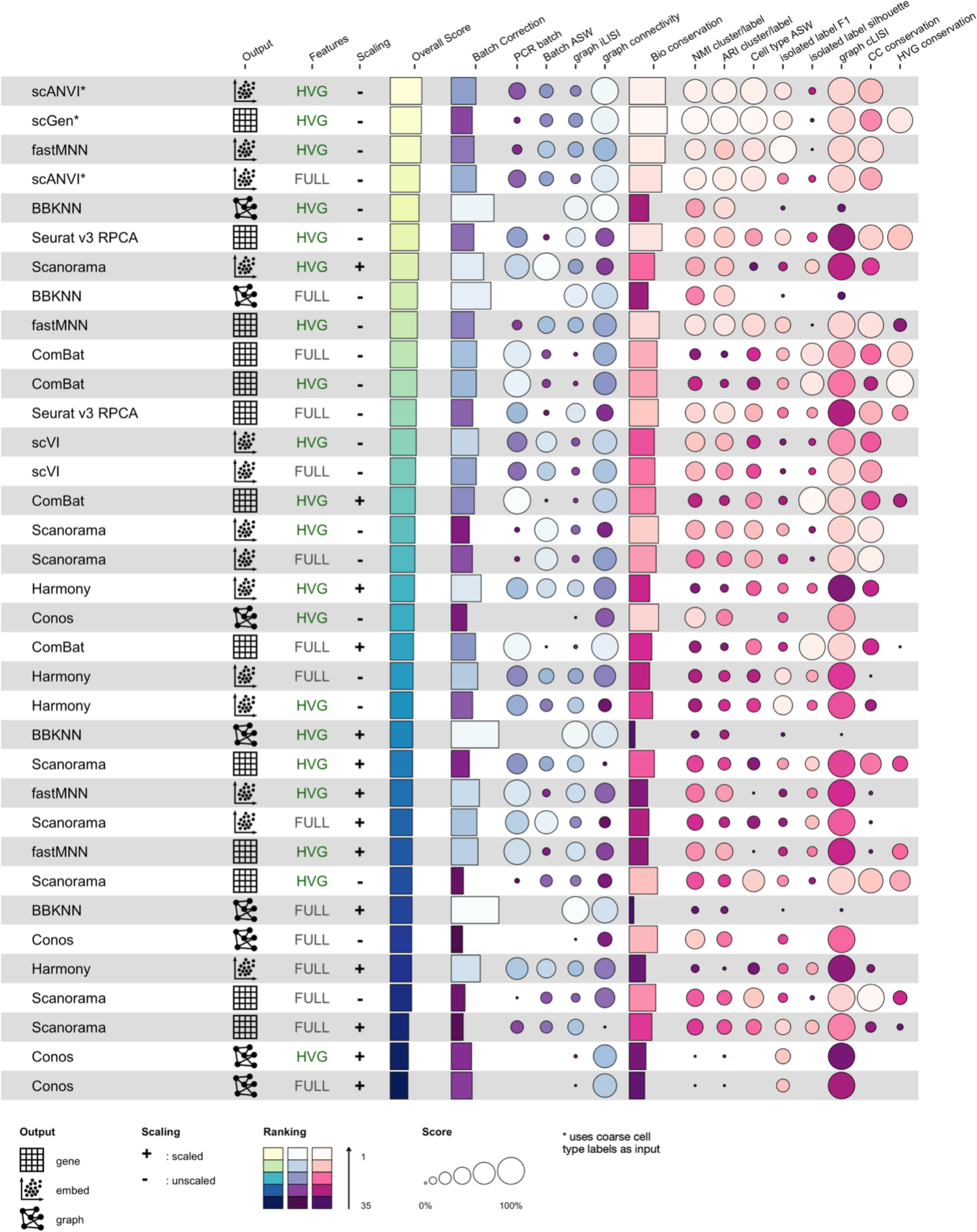
Results of dataset integration benchmarking. Rows represent methods tested, using a particular preprocessing. Preprocessing is summarized by “Features” (either HVG or FULL, corresponding to a gene selection of the 2000 or 6000 most highly variable genes, respectively) and “Scaling” (specifying whether or not gene values were scaled to mean 0 and standard deviation 1 across cells). Methods are sorted by overall score. Overall score is a weighted mean of the batch correction score and an th bio conservation score, which in turn are a mean of the individual metrics within the category. Metrics have been previously described^16^. The output column specifies whether a method has corrected gene counts, an integrated embedding, or an integrated graph as output. Scanorama and fastMNN were benchmarked twice, once with a corrected gene matrix as output, and once with a corrected embedding as output. scANVI and scGen were given coarse cell type labels as input, as indicated by the asterisk. fastMNN required too much memory on the full feature (6000 gene) data, therefore only the hvg results are included in the figure.

**Extended Data Figure 2.**
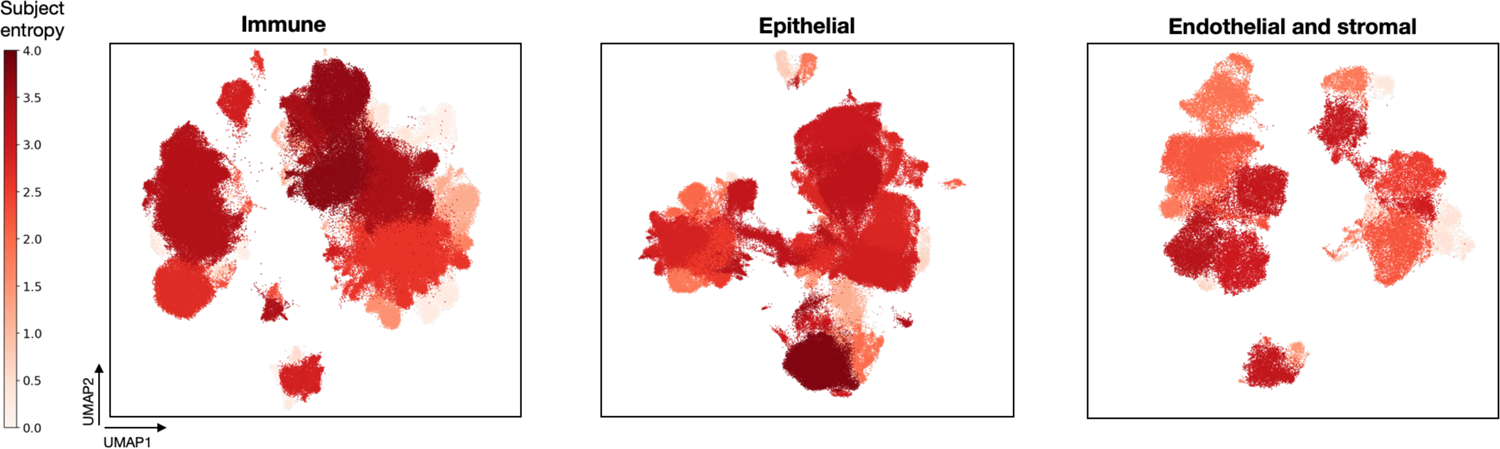
Subject diversity per HLCA core cluster. Subject diversity is calculated for every cluster as entropy of subject proportions in the cluster, with high entropy indicating the cluster contains cells from many different subjects. Matching cell type annotations are shown in fig. 3d.

**Extended Data Figure 3.**
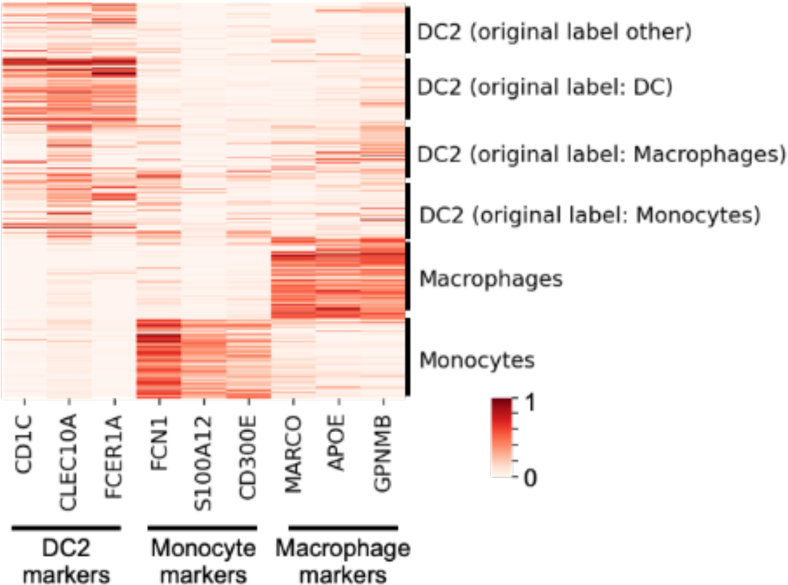
Marker expression among cells from a high-label-entropy cluster. DC2, monocyte and macrophage marker expression is shown for cells from the immune cluster with highest label entropy, as depicted in fig. 3c. Cells are labeled by their final annotation, as well as their original label. Log-normalized counts are scaled such that for each gene the 99th expression percentile, as calculated among all cells included in the heatmap, is set to 1.

**Extended Data Figure 4.**
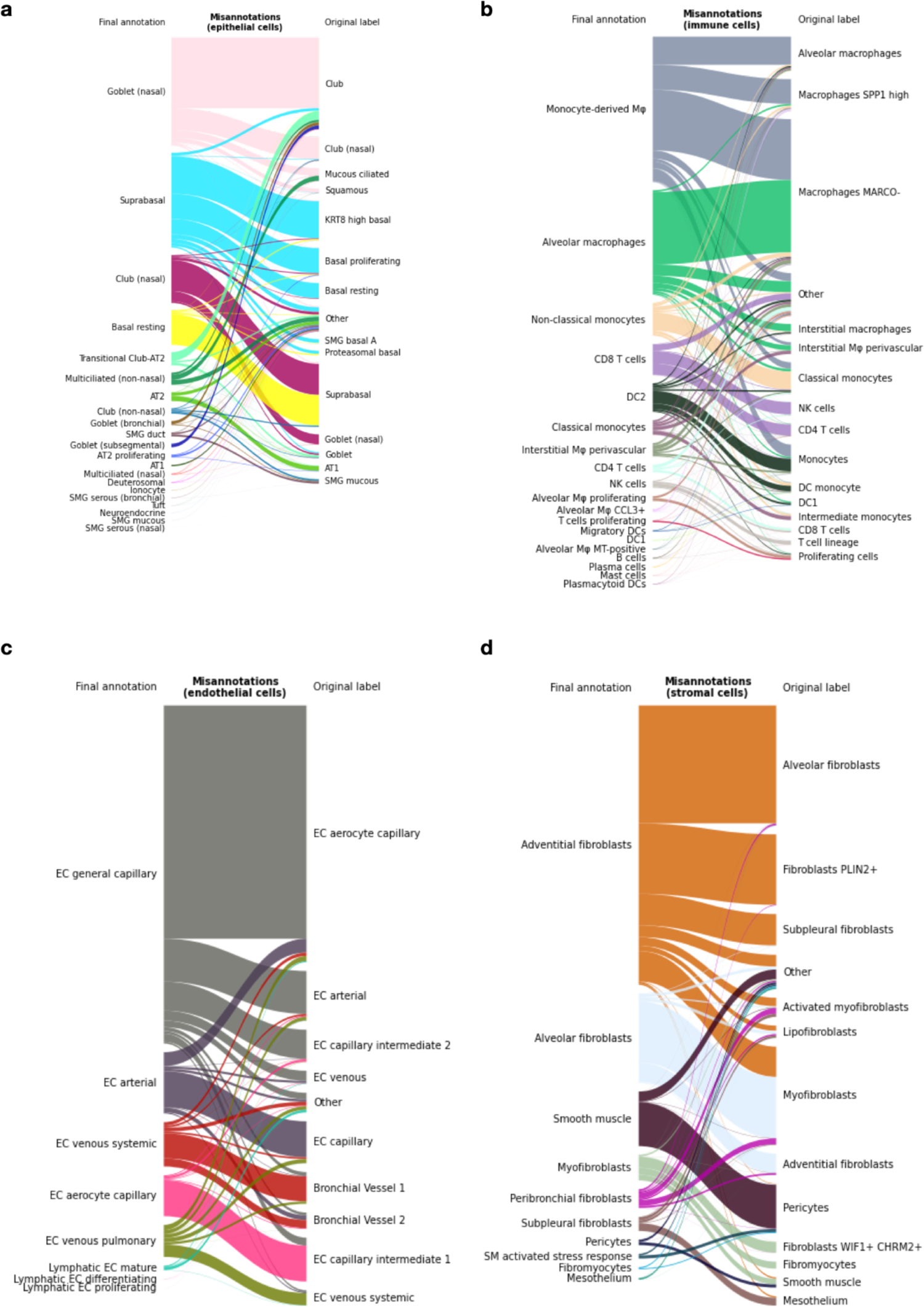
Final annotations and original annotations of mislabeled cells. **a**, Final annotations (left) and original harmonized labels (right) of epithelial cells for which the final annotation and original label are contradictory, i.e. misannotated cells. Original labels that represent less than 1% of epithelial cells are set to “Other”. **b, c, d**, as **a** but for immune, endothelial and stromal cells, respectively.

**Extended Data Figure 5.**
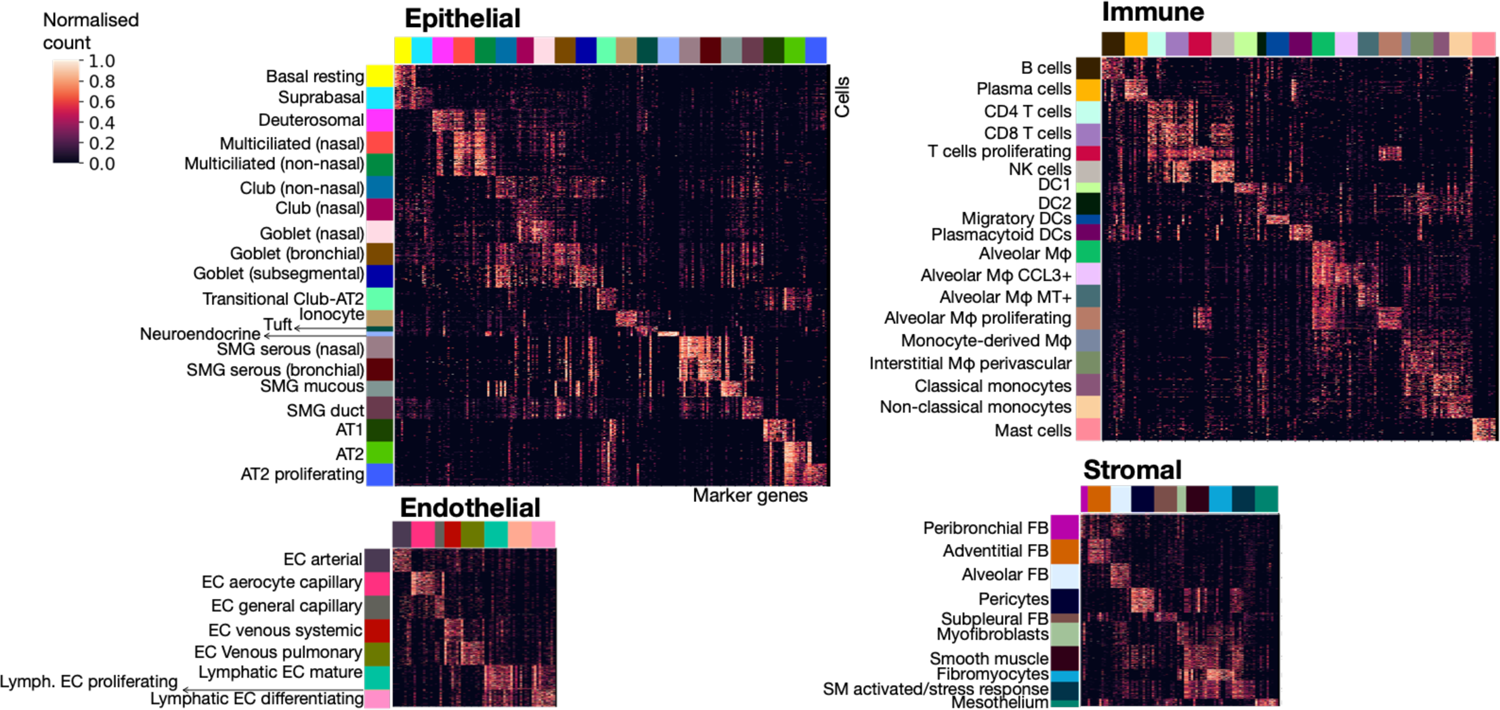
Expression of top 10 marker genes for every HLCA cell type annotation. Marker gene names are included in **Supplementary Data Table 5**. Where more than 750 cells of the cell type were present in the HLCA, expression is only shown for 750 randomly sampled cells from that cell type. Counts were normalized such that a gene’s 99th expression percentile across all cells for the heatmaps was set to 1. DC: dendritic cell. AT1, 2: alveolar type 1, 2 cells. EC: endothelial cells. FB: fibroblasts. Mφ: macrophages. MT: metallothionein. NK cells: natural killer cells. SM: smooth muscle. SMG: submucosal gland.

**Extended Data Figure 6.**
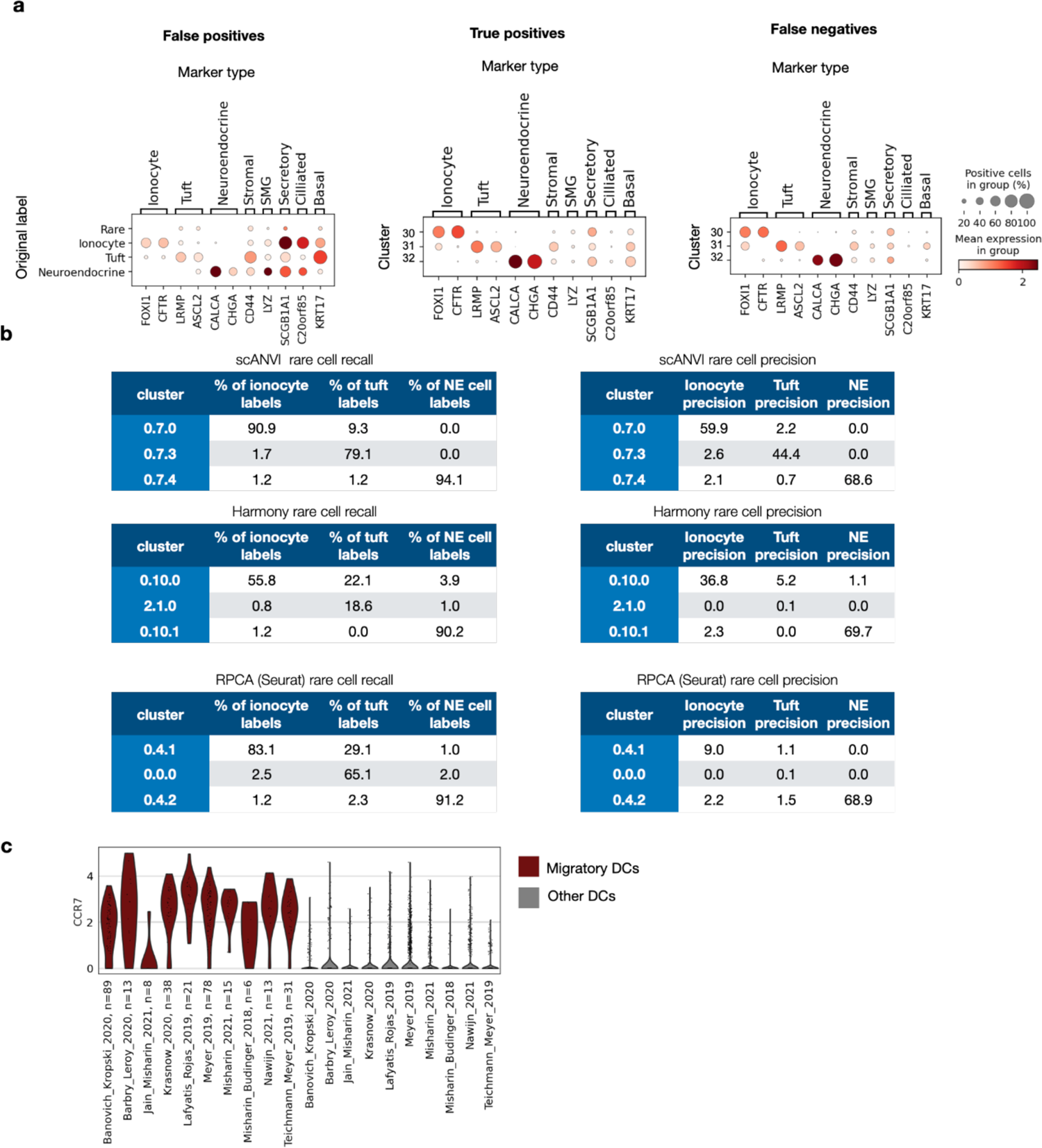
Rare cell identification and recovery for three different integration methods. **a**, Rare cell marker expression among different cell groups in the HLCA core. Markers of other cell types are included to show possible mis-annotations. Left: marker expression of cells originally labeled as rare, but who did not fall in one of the three rare cell clusters of the HLCA (“false positives”), subdivided by original label. Middle: Marker expression of cells originally labeled as rare, and who fell in one of the three rare cell clusters in the HLCA (“true positives”), subdivided by cluster. Right: marker expression of cells who were not originally labeled as rare, but which nonetheless were classified in one of the three rare cell clusters of the HLCA core (“false negatives”), subdivided by cluster. **b**, Recall and precision of rare cell types in distinct clusters for three different integration methods: scANVI^24^, Harmony^32^ and Seurat’s RPCA^28^. Results are shown for the best-performing preprocessing for each method, and based on the benchmarking data (12 datasets from the HLCA core). Recall (i.e. the percentage of cells with a specific label that are present in the cluster under consideration) and precision (i.e. the percentage of cells from a cluster labeled as the cell type under consideration) are shown for the three level 3 clusters with the highest recall of ionocytes, tuft, and NE cells respectively. NE: neuroendocrine. **c**, Expression of migratory DC marker *CCR7* among migratory DCs versus other DCs, split by study. Number of migratory DCs per study is specified in the x-axis labels. DC: dendritic cell.

**Extended Data Figure 7.**
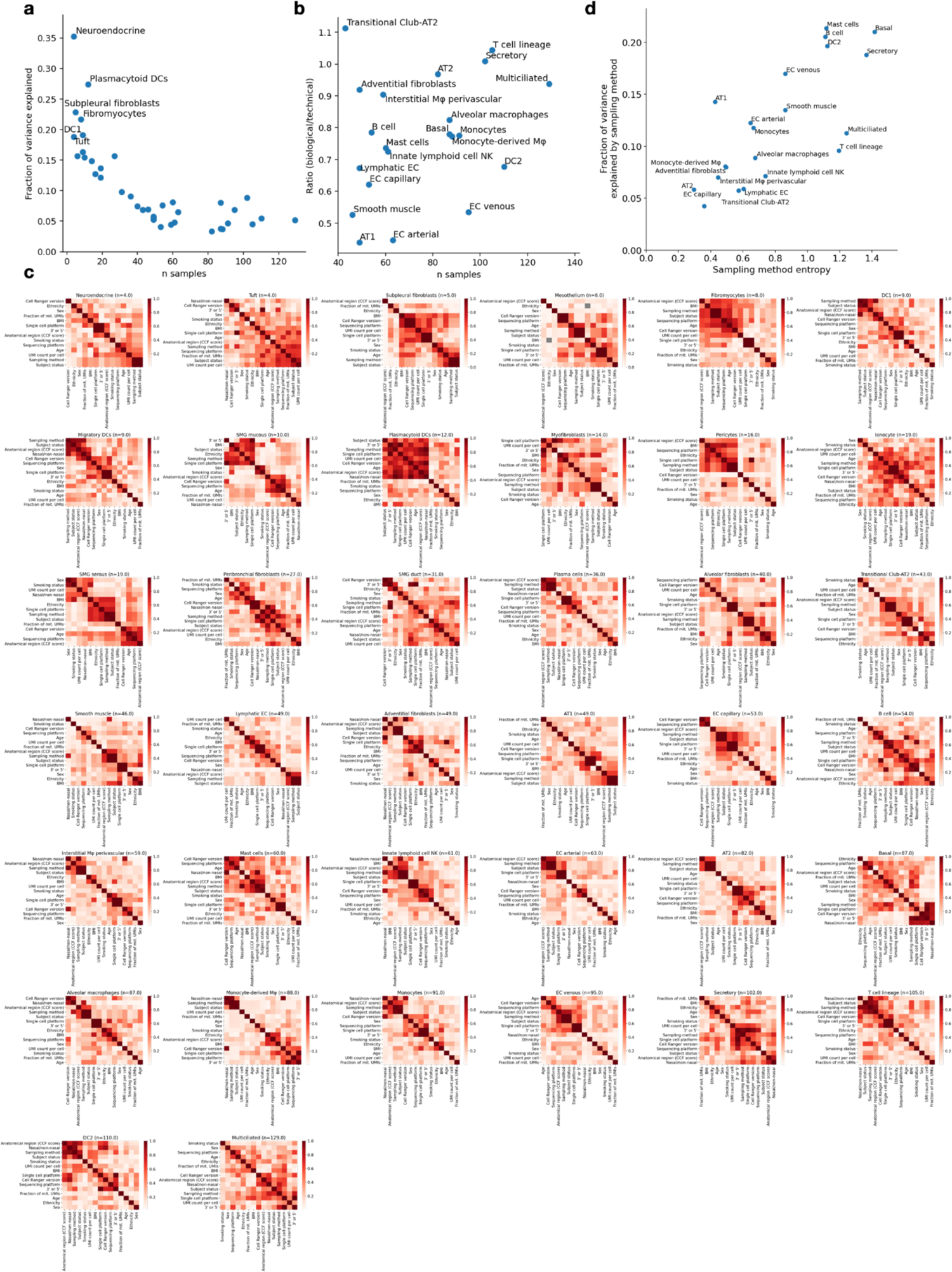
Correlation of technical and biological covariates with number of samples, and with each other. **a**, Relation between the number of samples in which a cell type was observed, and the mean inter-sample variance explained by the technical and biological covariates. After n=40, these two variables become independent. Inter-sample variance was calculated based on the scANVI-integrated embedding, taking the mean score of each latent dimension across cells, for every sample for every cell type. **b**, The ratio of mean variance explained by biological covariates over that explained by technical covariates for different cell types (y-axis), and its association with the number of samples in which a cell type was observed (x-axis) (Pearson r: 0.28, p=0.21). **c,** Correlation between covariates for every cell type, calculated at sample level. The square root of the fraction of variance from one covariate that could be explained by the other covariate through linear regression is shown (equivalent of Pearson r for two continuous covariates). If both covariates were categorical and had more than two categories, normalized mutual information was calculated instead. **d,** Variance explained by sampling method depends only partly on diversity in sampling methods as quantified by the sampling method entropy (Pearson r=0.75, p=0.0001). Some cell types, such as T cells, are less sensitive to sampling than others, despite higher sampling method entropy. AT1, 2: alveolar type 1, 2 cells. DC: dendritic cells. DC1, 2: DC type 1, 2. EC: endothelial cells. Mφ: macrophages. NK cells: natural killer cells. NKT cells: natural killer T cells. SMG: submucosal gland.

**Extended Data Figure 8.**
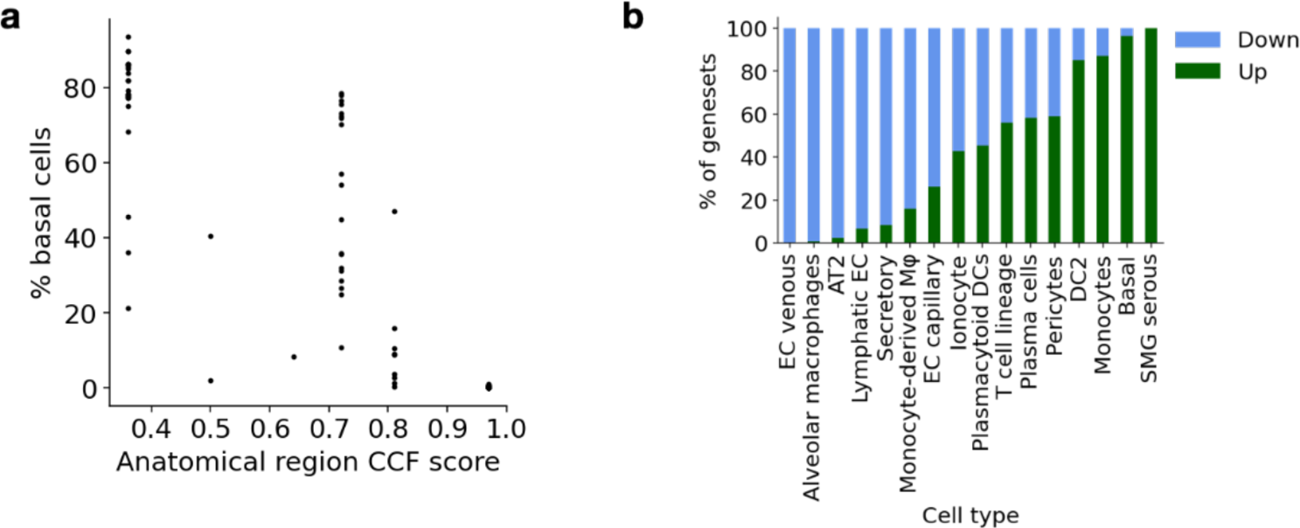
Trends associated with BMI and sample anatomical location. **a**, The percentage of basal cells per sample decreases with higher anatomical region CCF score, i.e. towards the distal lung (Spearman r=-0.89, p<0.001). CCF: common coordinate framework. **b**, Proportions of gene sets significantly up- and down-regulated with BMI, per cell type. AT2: Alveolar type 2 cells. DC: dendritic cells. DC2: DC type 2. EC: endothelial cells. Mφ: macrophages. SMG: submucosal gland.

**Extended Data Figure 9.**
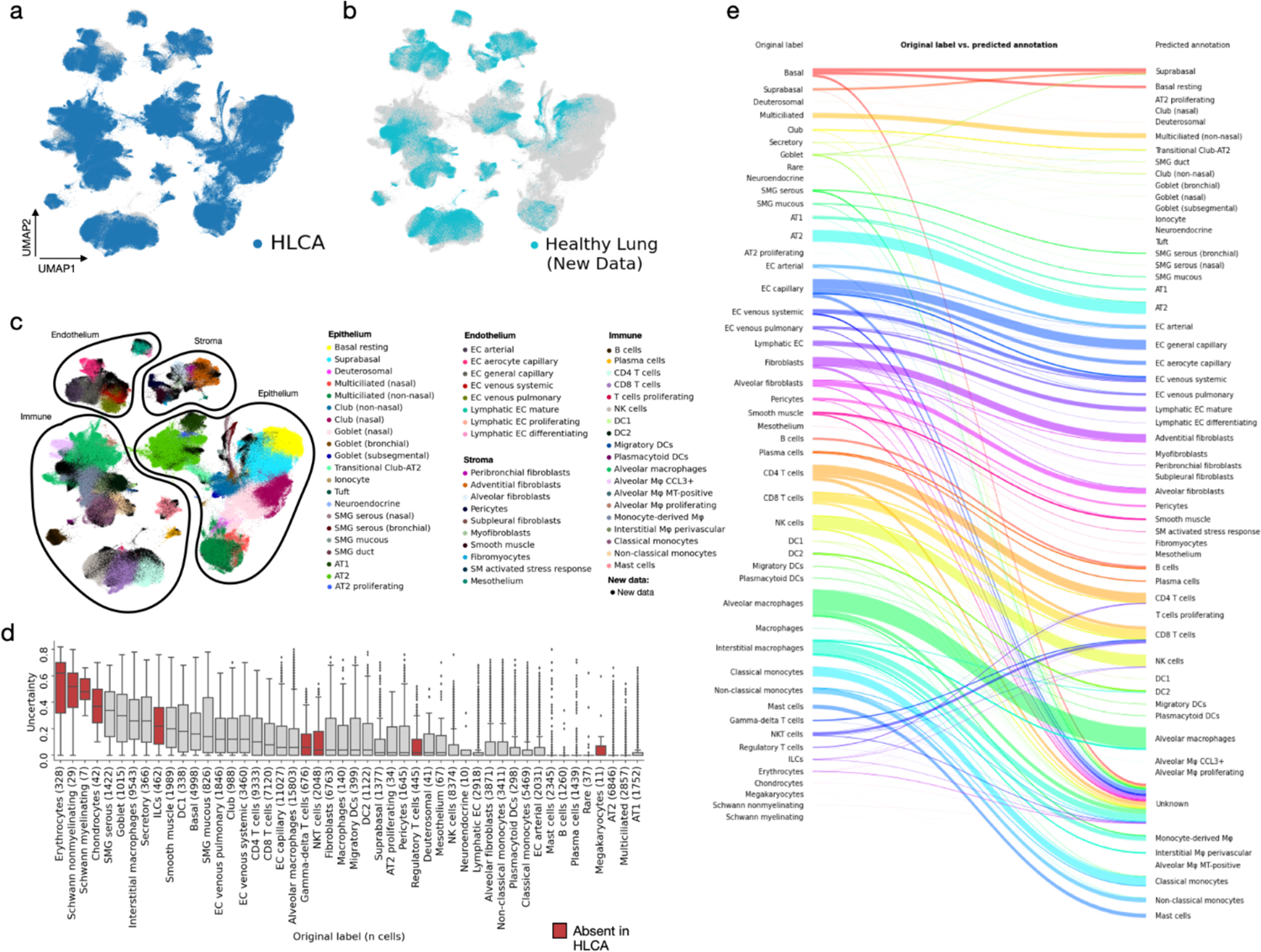
Mapping of unseen healthy lung scRNA-seq data to the HLCA core. **a**, A UMAP of the jointly embedded HLCA core and the newly mapped healthy lung data. Cells from the new data are shown in gray, cells from the HLCA are shown in blue and plotted on top. **b**, Same as a, but now plotting cells from the HLCA in gray, and cells from the new data on top in light blue. **c**, Same as a, but now coloring cells from the HLCA core by their final annotation, and coloring cells from the new data in black. **d**, Uncertainty of label transfer (ranging from 0 to 1) for cells from the mapped data, subdivided by original cell type label. Number of cells per label is shown between brackets. Cell labels are ordered by mean uncertainty. Boxes of cell labels not present in the HLCA core are colored red. Boxes show median and interquartile range of uncertainty. Cells with uncertainties more than 1.5 times the interquartile range away from the high and low quartile are considered outliers and plotted as points. Whiskers extend to the furthest non-outlier point. **e**, Sankey plot of original labels of cells from the mapped dataset versus predicted annotations based on label transfer. Cells with uncertainty >0.3 are labeled “unknown”. AT1, 2: alveolar type 1, 2 cells. DC: dendritic cells. DC1, 2: DC type 1, 2. EC: endothelial cells. ILCs: innate lymphoid cells. MT: metallothionein. Mφ: macrophages. NK cells: natural killer cells. NKT cells: natural killer T cells. SM: smooth muscle. SMG: submucosal gland.

**Extended Data Figure 10.**
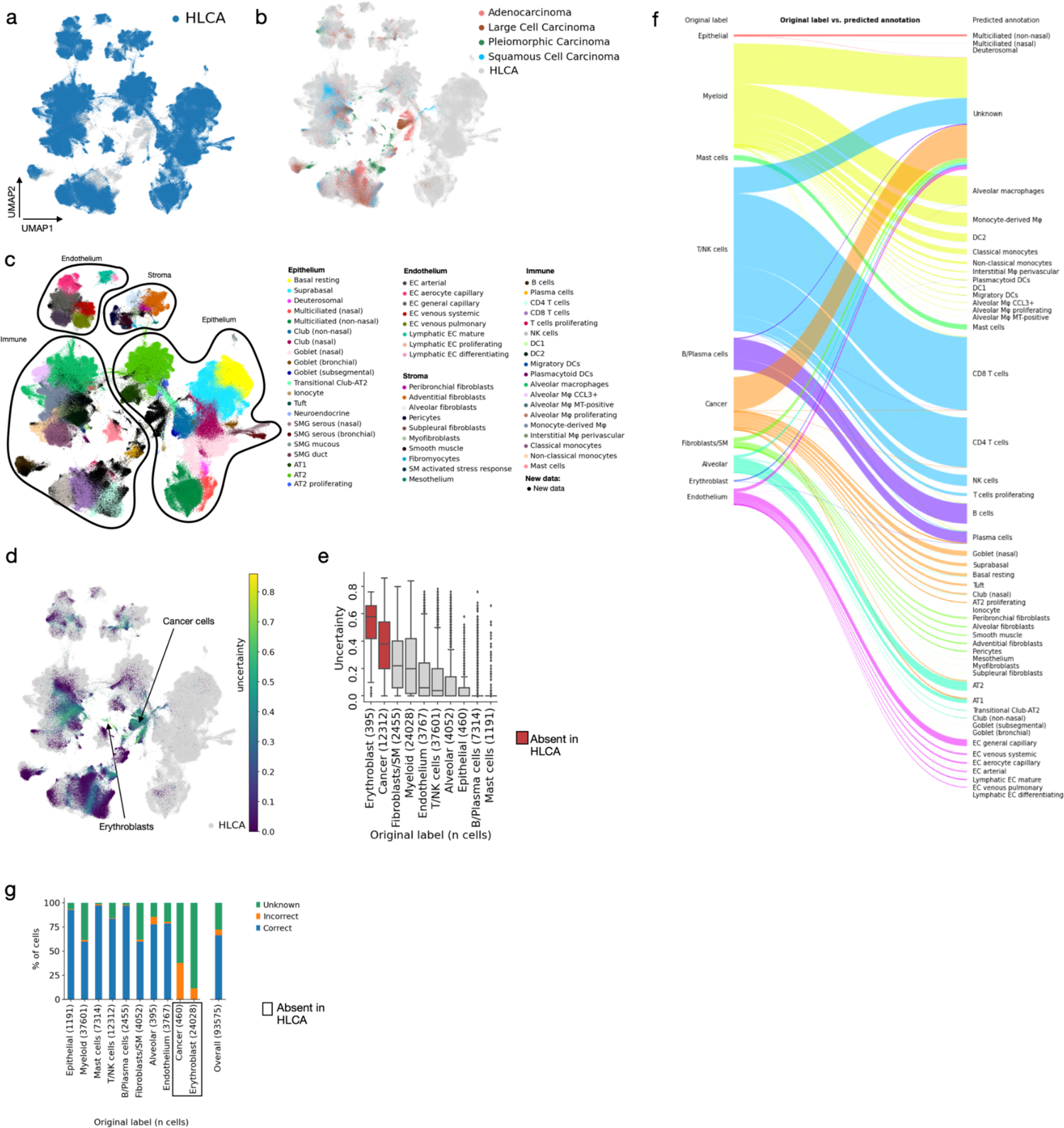
Mapping of unseen lung cancer data to the HLCA. **a**, A UMAP of the jointly embedded HLCA and lung cancer data. Cells from the lung cancer dataset are shown in gray, cells from the HLCA core are shown in blue and plotted on top. **b**, Same as a, but now plotting cells from the HLCA core in gray. Cells from the mapped data are plotted on top, and colored by the cancer type of the patient. **c**, Same as a, but now coloring cells from the HLCA core by their final (coarse) annotation, and coloring cells from the mapped cancer data in black. **d**, Uncertainty of label transfer, shown for all cells from the mapped data. Regions dominated by high-uncertainty cells are labeled by the original cell type label. **e**, Uncertainty of label transfer (ranging from 0 to 1) for the mapped cells, subdivided by original cell type label. Number of cells per label is shown between brackets. Boxes of cell type labels not present in the HLCA core are colored red. Cell types are ordered by mean uncertainty. Boxes show median and interquartile range of uncertainty. Cells with uncertainties more than 1.5 times the interquartile range away from the high and low quartile are considered outliers and plotted as points. Whiskers extend to the furthest non-outlier point. **f**, Sankey plot of original labels of the mapped data versus predicted annotations based on label transfer. Cells with uncertainty >0.3 are labeled “unknown”. **g**, Percentage of cells from newly mapped healthy lung dataset that are either annotated correctly or incorrectly by label transfer annotation, or annotated as unknown, subdivided by original cell type label. The number of cells in the mapped dataset labeled with each label are shown between brackets after cell type names. Cell type labels not present in the HLCA are boxed. AT1, 2: alveolar type 1, 2 cells. DC: dendritic cells. DC1, 2: DC type 1, 2. EC: endothelial cells. MT: metallothionein. Mφ: macrophages. NK cells: natural killer cells. NKT cells: natural killer T cells. SM: smooth muscle. SMG: submucosal gland.

**Extended Data Figure 11.**
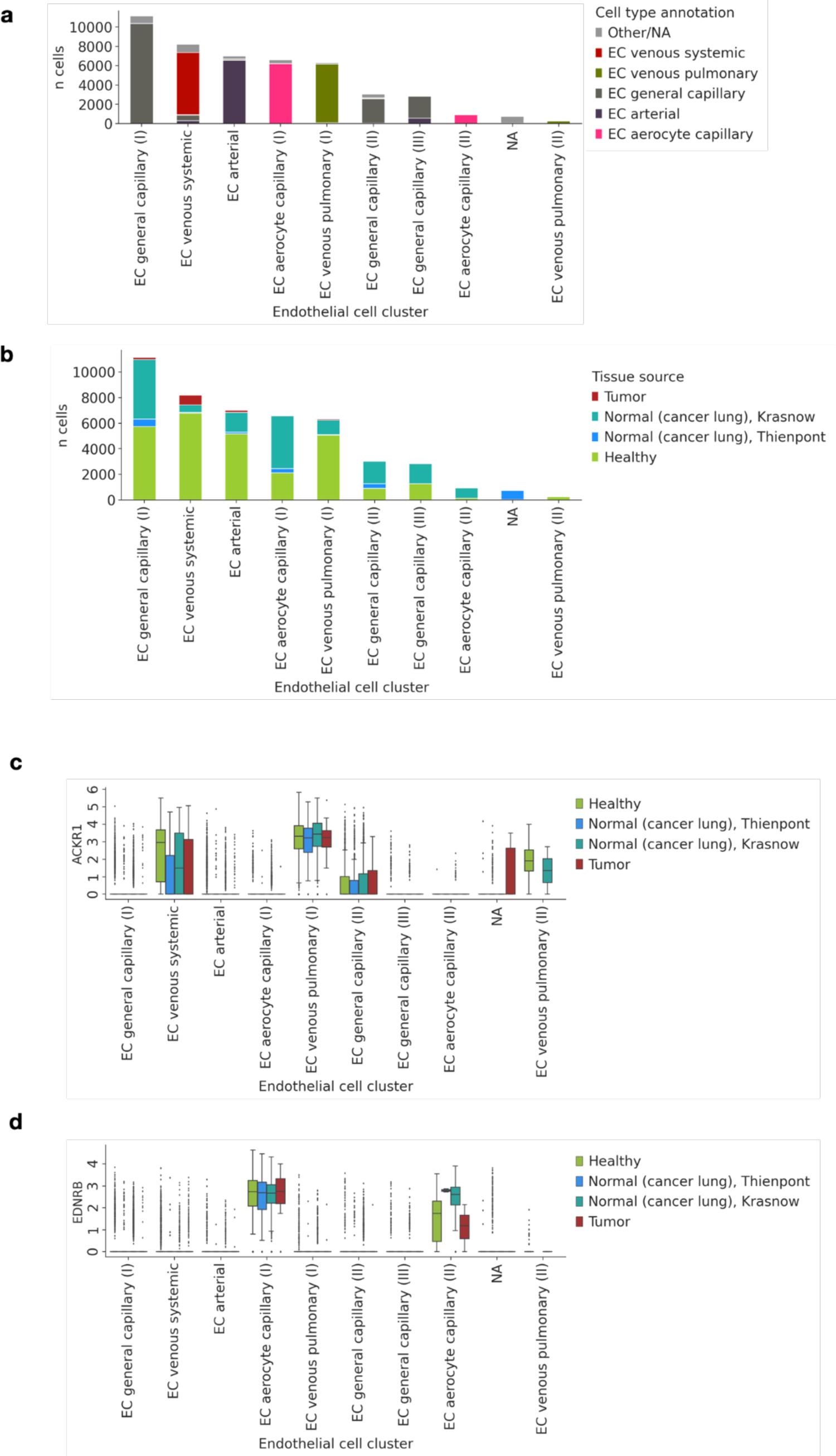
Endothelial cell clustering and marker expression in the joint embedding of unseen lung cancer data and the HLCA core. **a**, Cell type composition of EC clusters from the joint embedding of the HLCA core and the mapped lung cancer data. Final HLCA core annotations are shown, with cells from the cancer data as well as HLCA core annotations with fewer than 50 cells set to “Other/NA”. Clusters are named by their dominant cell type. b, Tissue source composition of EC clusters. Tissue source is either tumor, normal (but from cancer patients), or healthy (from subjects without lung cancer). Normal is split by dataset, including the Krasnow dataset from the HLCA core (with non-tumorous tissue from lung cancer patients) for comparison. c, ACKR1 expression in EC clusters, split by tissue source. Boxes show median and interquartile range of expression. Cells with uncertainties more than 1.5 times the interquartile range away from the high and low quartile are considered outliers and plotted as points. Whiskers extend to the furthest non-outlier point. d, same as c, but now showing *EDNRB* expression. EC: endothelial cell.

**Extended Data Figure 12.**
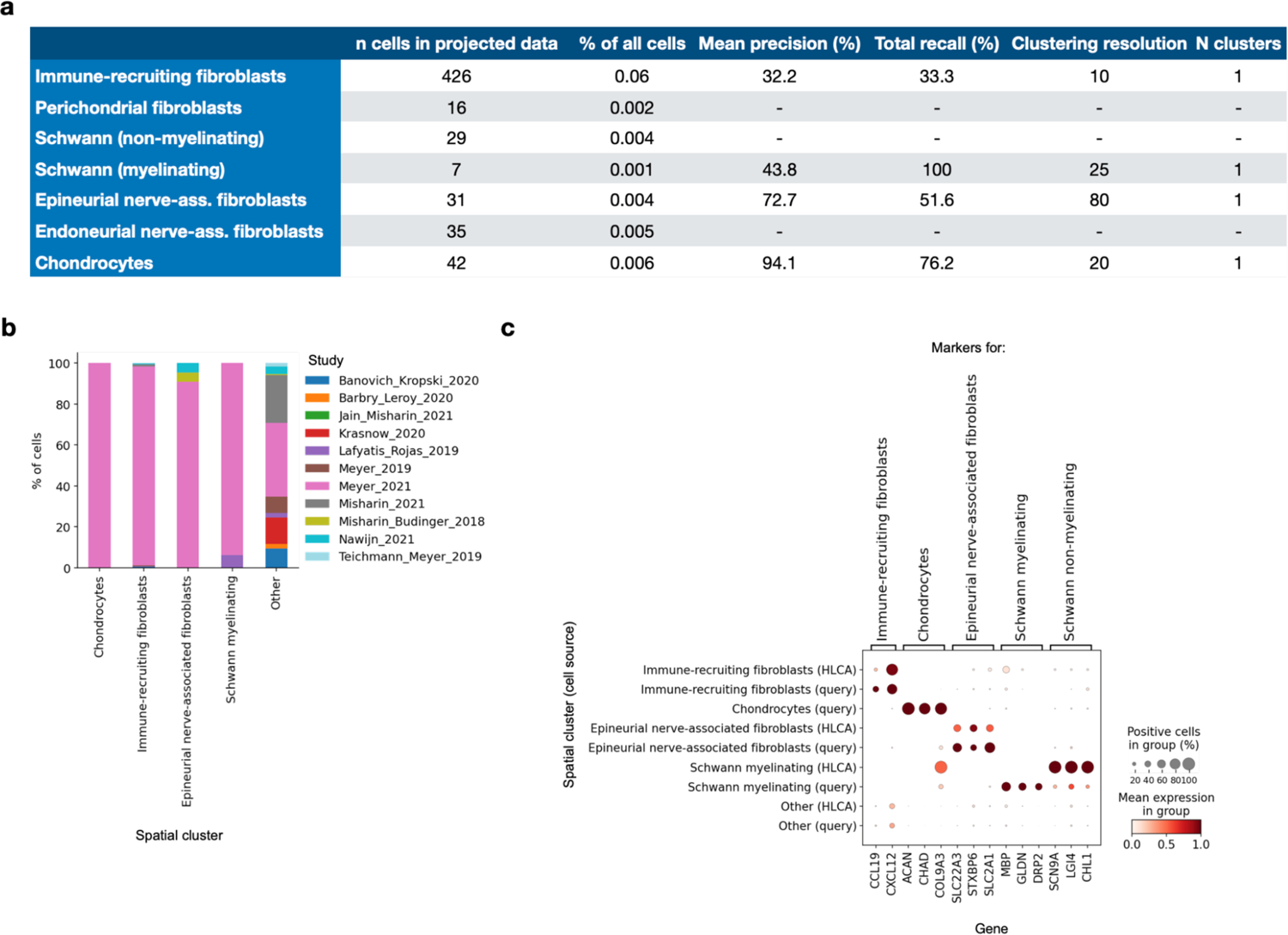
Identification of spatial-location-based cell types in the HLCA core based on mapping of spatially labeled data. **a**, Precision (i.e. the percentage of cells from a cluster labeled as the cell type under consideration) and recall (i.e. the percentage of cells with a specific label that are present in the cluster under consideration) of spatial cell types in spatial clusters. Precision and recall were calculated among cells of the mapped data only. Clustering resolution at which the cluster was identified, and number of clusters per spatial cell type is also shown. % of all cells specifies the percentage of cells with the label among cells of both the projected data and the HLCA core. b, Composition of spatially annotated clusters in terms of study from which the cells came. c, Marker expression of spatially annotated cell type markers across spatially annotated clusters, splitting clusters in cells from the HLCA core, and cells from the newly mapped data (“query”). Gene expression was normalized to range, within fibroblasts, from 0 to 1.

**Extended Data Figure 13.**
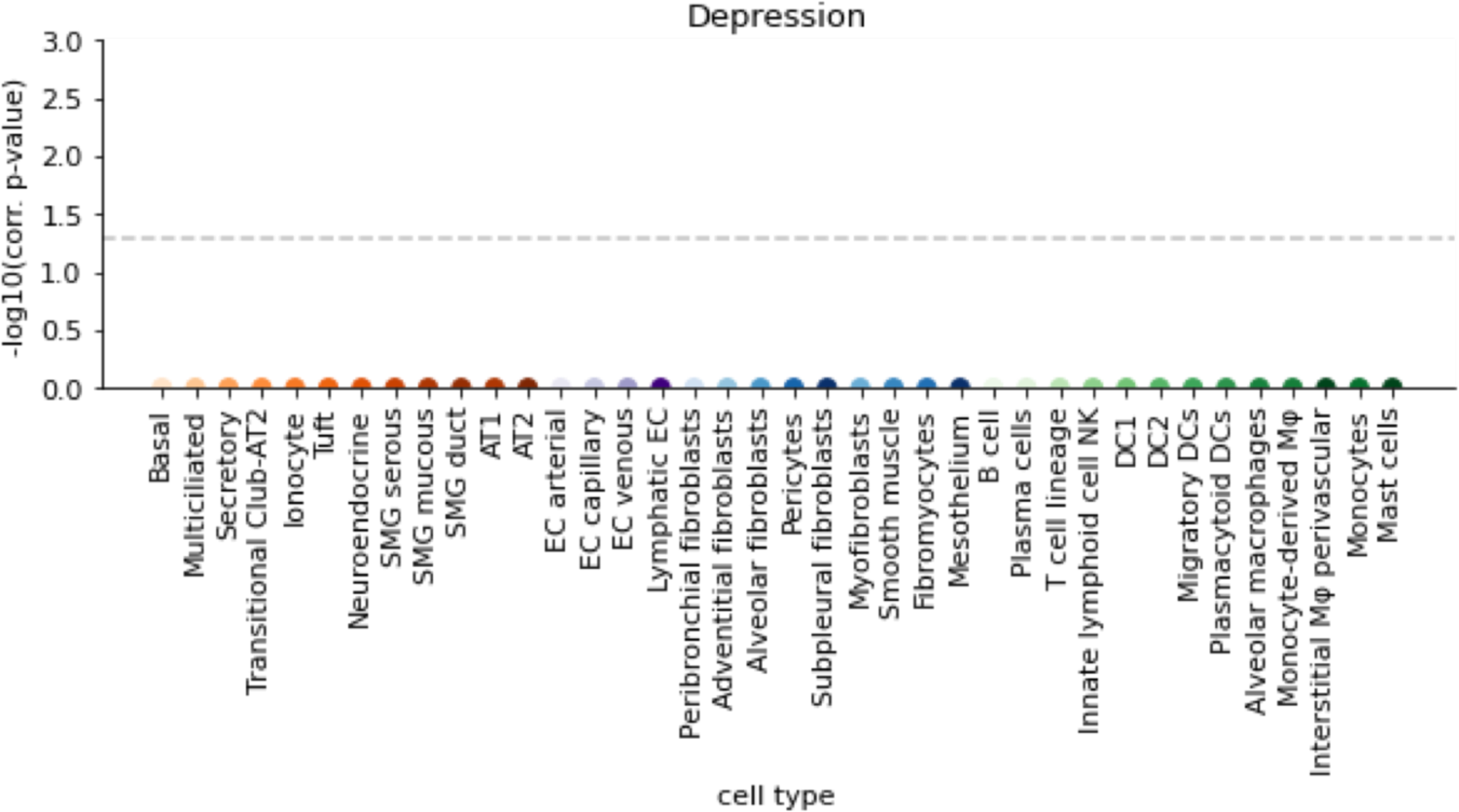
Association of lung cell types with depression. Negative control for analysis of fig. 5d. Association is based on phenotype-related genomic variants from a GWAS study on depression, and cell-type specific differentially expressed genes calculated from the HLCA core. Horizontal dashed line indicates significance threshold alpha=0.05. p-values are multiple-testing-corrected with the Benjamini-Hochberg procedure.

**Extended Data Figure 14.**
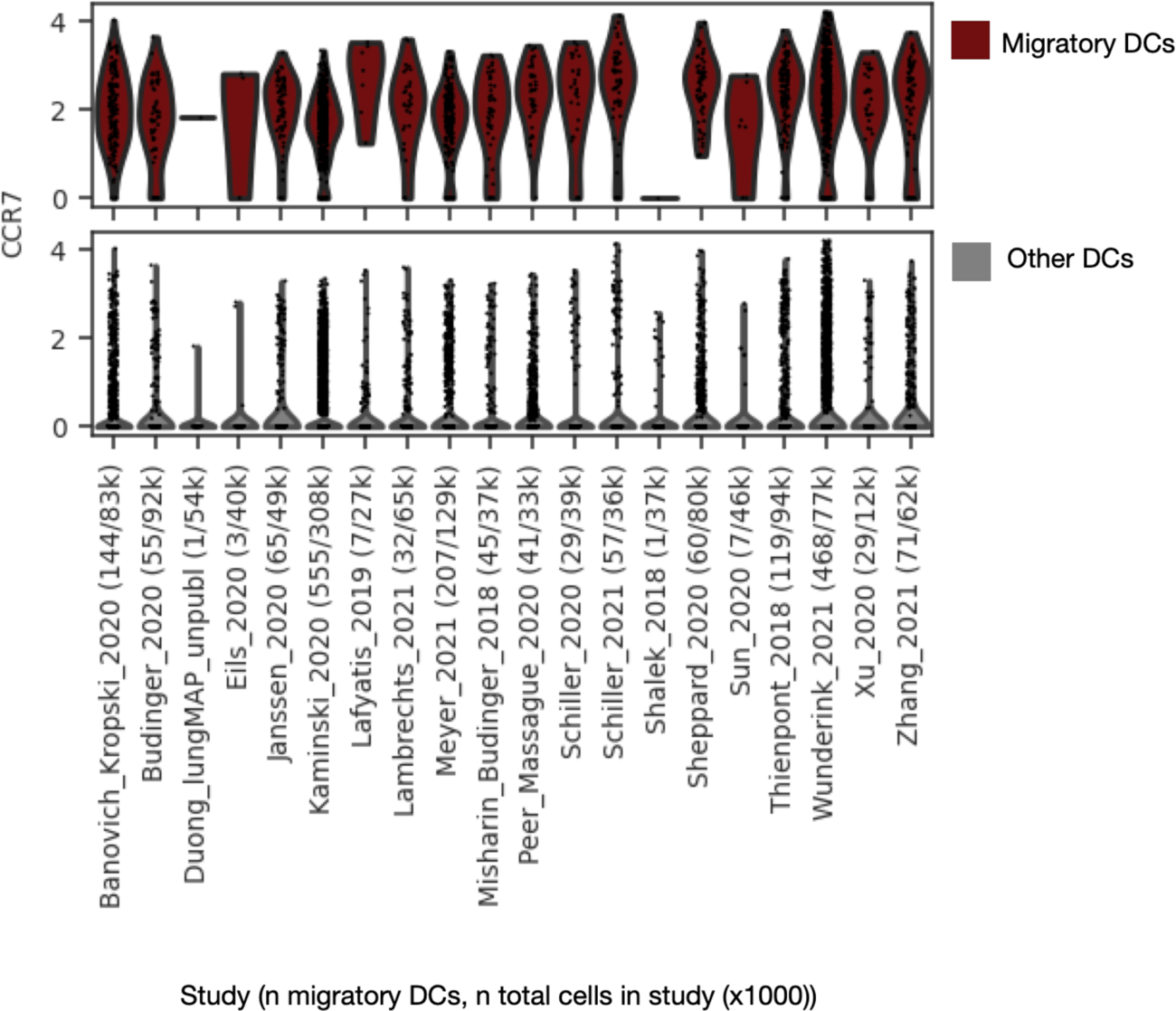
Expression of *CCR7* among cells annotated as migratory DCs by label transfer. Expression of *CCR7* is shown for all cells that were annotated as migratory DCs with low uncertainty (<0.2) (top) and all other cells annotated as DC (bottom) by label transfer from the HLCA core to the extended HLCA. Cells are grouped based on study of origin (some studies contain multiple datasets). X-tick labels show study, number of cells annotated as migratory DCs, and number of total cells (in thousands) per study. *CCR7* counts shown are counts that were normalized based on the total count among 2000 genes used for mapping to the HLCA core, and then log-transformed. DCs: dendritic cells.

**Extended Data Figure 15.**
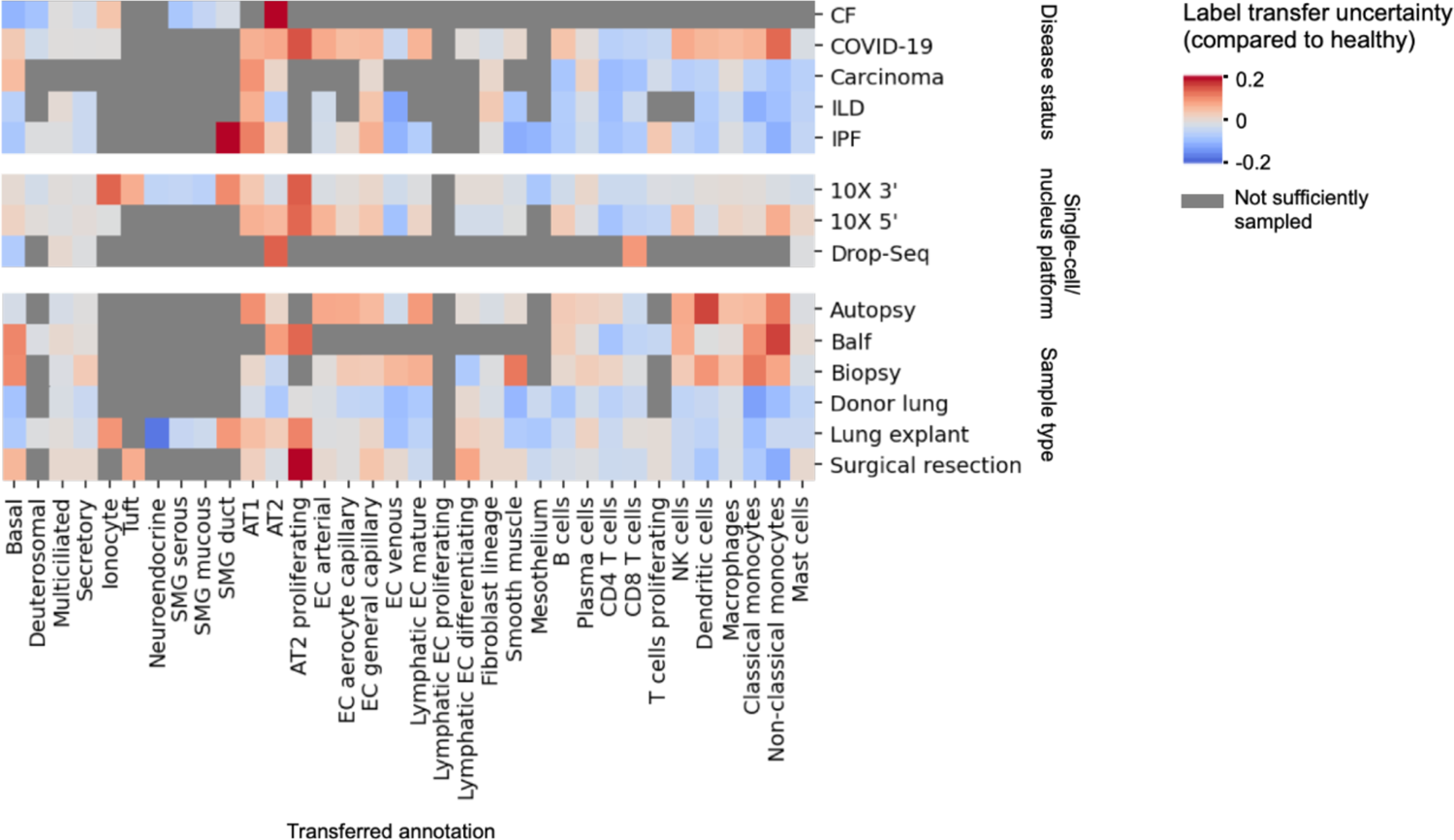
Label transfer uncertainty per cell type across different experimental features. Label transfer uncertainty of cell types is shown for categories of three experimental features, as compared to uncertainty in healthy cells. For every category and for each cell type, the mean uncertainty across datasets from that category was calculated, using per-dataset means and splitting up datasets with samples from more than one category. The difference between cell type uncertainty from each category and those of healthy datasets is shown. Where mean uncertainty was higher than 0.25 even in healthy, coarser parent labels were included (e.g. for macrophages) instead of the finest cell type annotations. Datasets with fewer than 20 cells of a cell type are excluded for that cell type. When no dataset sampled enough cell types, the plot is masked in gray. Values higher than 0.2 or lower than −0.2 are cut off to 0.2 and −0.2, respectively.

**Extended Data Figure 16.**
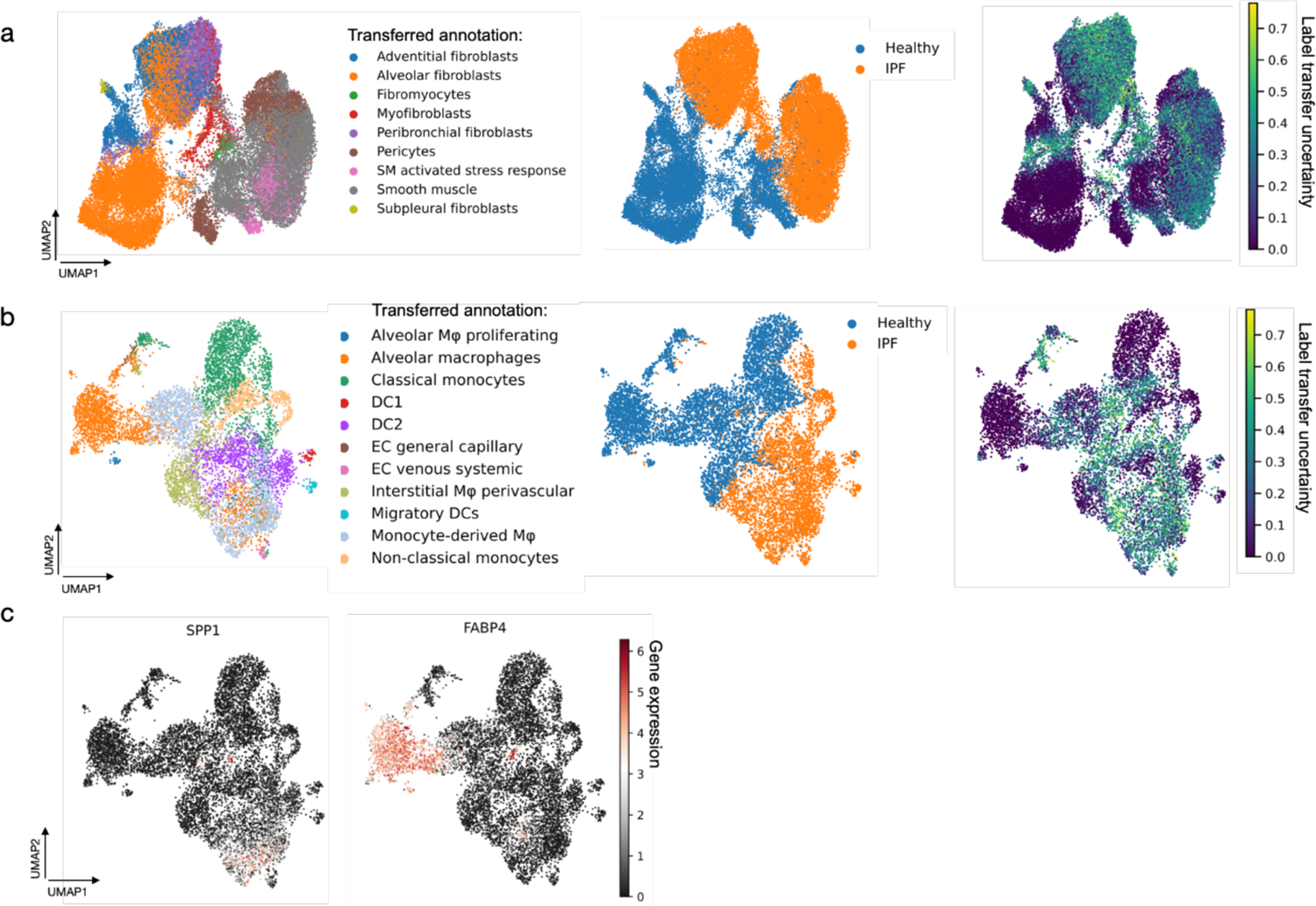
Transferred labels and matching uncertainty for a mapped IPF dataset. **a**, UMAPs of cells originally labeled as stroma, from a mapped IPF dataset^62^ including both healthy and IPF samples. Cells are labeled by annotation transferred from the HLCA core (left), by disease status (middle), and by label transfer uncertainty (right). Cells with labels transferred to fewer than 10 cells were excluded. **b**, same as a, but showing cells originally labeled as macrophages. **c**, As b, but now colored by expression of *SPP1* and *FABP4*.

**Extended Data Figure 17.**
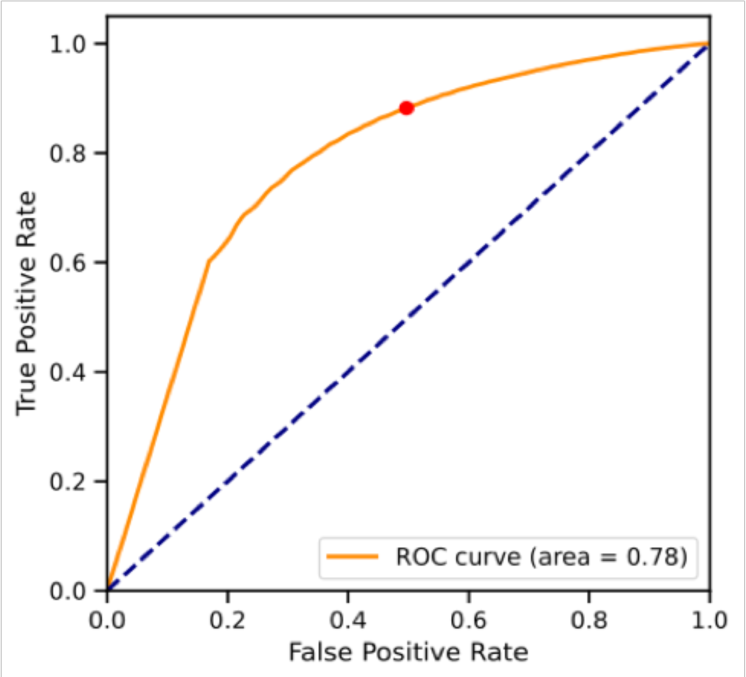
Calibration of label transfer uncertainty cutoff. ROC curve of label transfer accuracy across 7 datasets. The true and false positive rate of the chosen cutoff point (0.2), below which transferred labels will be considered low uncertainty, are shown as a red point on the ROC curve.

## Notes

https://beta.fastgenomics.org/p/hlca

https://cellxgene.cziscience.com/collections/6f6d381a-7701-4781-935c-db10d30de293

